# FocalSV: target region-based structural variant assembly and refinement using single-molecule long read sequencing data

**DOI:** 10.1101/2024.11.21.624735

**Authors:** Can Luo, Zimeng Zhou, Xin Maizie Zhou

**Author notes:** Equal contributor (Can Luo, Zimeng Zhou).

## Abstract

Structural variants (SVs) play a critical role in shaping the diversity of the human genome and their detection holds significant potential for advancing precision medicine. Despite notable progress in single-molecule long-read sequencing technologies, accurately identifying SV breakpoints and resolving their sequence remains a major challenge. Current alignment-based tools often struggle with precise breakpoint detection and sequence characterization, while whole genome assembly-based methods are computationally demanding and less practical for targeted analyses. Neither approach is ideally suited for scenarios where regions of interest are predefined and require precise SV characterization. To address this gap, we introduce FocalSV, a target region assembly-based SV detection tool that combines the precision of assembly-based methods with the efficiency of region-specific approaches. FocalSV was evaluated on nine germline datasets and two paired normal-tumor cancer datasets, demonstrating superior performance in both precision and efficiency.

## Background

Single nucleotide variants (SNVs) constitute the most abundant form of genetic variation in humans and can be efficiently detected using short-read sequencing technologies. Therefore, genome-wide association studies (GWAS) have primarily focused on SNVs to investigate the genetic basis of phenotypic traits. In contrast, structural variants (SVs) - larger genomic alterations of 50 base pairs (bp) or more, including insertions (INS), deletions (DEL), duplications (DUP), inversions (INV), and translocations (TRA) - represent a major source of genetic diversity but are more challenging to detect accurately with short-read sequencing [1]. Despite their considerable biomedical relevance [2], short-read-based approaches fail to identify over half of the SVs within an individual genome [3], constraining our ability to fully delineate the genetic landscape associated with complex disease phenotypes.

In recent years, long-read sequencing technologies have significantly advanced SV detection by providing extended read lengths, making it possible to resolve complex genomic regions that were previously challenging to analyze [4, 5]. Technologies such as PacBio’s HiFi and Continuous Long Reads (CLR) and Oxford Nanopore Technology (ONT) offer distinct advantages. PacBio’s HiFi reads, for instance, achieve accuracy of up to 99.9% with read lengths up to 20 kb, making them particularly well-suited for capturing SVs in complex genomic contexts [6]. ONT, on the other hand, can generate even longer reads - exceeding 1 megabase on some cases - though with slightly lower accuracy. These long-read technologies allow for more accurate genomic alignment and improved assembly, significantly enhancing confidence in SV detection and enabling a more comprehensive view of the genomic landscape [7]. However, despite these advantages, long-read technologies come with notable limitations. High-quality long reads remain costly and require substantial computational resources for processing, particularly when performing whole-genome assembly. These challenges present significant barriers in large-scale studies, where analyzing SVs across entire genomes for hundreds to thousands of samples is computationally intensive and time-consuming. Consequently, there is increasing demand for more efficient approaches that leverage long-read capabilities without the high computational cost of whole-genome assembly.

The alternative approach, alignment-based methods, detects SVs by mapping long reads directly to a reference genome and identifying breakpoints based on alignment discrepancies. These methods are generally less computationally intensive and are well-suited for large-scale datasets. However, they face challenges in accurately identifying SV breakpoints and sequences compared to assembly-based methods [8]. Furthermore, these limitations in both whole genome-based and alignment-based methods make them suboptimal for scenarios requiring precise SV characterization in predefined genomic regions.

In this study, to address this gap, we developed FocalSV, a target region assembly-based SV detection tool that combines the precision of assembly-based methods with the efficiency of a region-specific approach. The region-specific design of FocalSV is particularly valuable for clinical and genomic research, enabling users to focus on medically relevant SVs in specific loci or regions with SVs of interest. We benchmarked FocalSV against several state-of-the-art alignment-based and assembly-based tools, demonstrating overall superior accuracy and robustness across a wide variety of long-read datasets. FocalSV provides a practical and scalable alternative to whole-genome assembly, making it suitable for analyzing regions of interest in both individual samples and largescale population studies.

## Methods

We present FocalSV’s approach for region-based assembly and refinement of SVs across each type of SVs: insertions (INSs), deletions (DELs), duplications (DUPs), translocations (TRAs), and inversions (INVs).

### Large INS and DEL assembly

FocalSV utilizes the whole genome aligned reads BAM file and the specified target regions as inputs to perform large indel SV (≥50bp) assembly and refinement. The target region-based workflow for identifying a single target SV involves the following steps: (1) Extraction of region-specific BAM file, (2) partitioning of reads, (3) haplotype-aware local assembly, (4) detection of candidate SV, and (5) filtering of indel SVs and refinement of genotypes.

#### Extraction of region-specific BAM file

FocalSV is designed for situations where regions suspected of containing SVs have been identified, but the precise details of the SVs within that region have yet to be determined. FocalSV assists in more accurately inferring SV breakpoint and sequence based on prior knowledge. To use FocalSV, users must provide regions suspected SV-containing regions, which serve as input. FocalSV then extracts region-specific BAM files for subsequent SV assembly and refinement. In our experiments, to validate the accuracy of SV calling, we adopted the benchmark VCF file of the sample HG002 from Genome in a Bottle (GIAB) as prior knowledge. Specifically, for INSs, the start and end positions of the target regions we utilized are defined as follows:

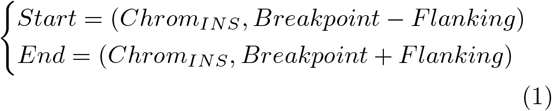

where the breakpoint refers to the insertion breakpoint on the reference genome, and the flanking size we used is 50kb by default.

In terms of DELs, the target regions we utilized are defined as follows:

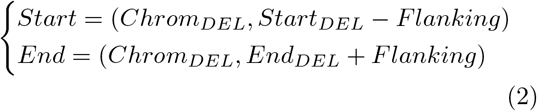

where the *start*_*DEL*_ and *end*_*DEL*_ refer to the starting and ending positions of a target DEL on the reference genome and the flanking size we used is 50kb. When the breakpoints of INSs and DELs are unknown, users can estimate breakpoints to define the target regions and adjust the flanking regions to ensure regions are sufficiently large to encompass the potential SV.

#### Partitioning of reads

The FocalSV pipeline integrates Longshot [9], a haplotype estimation tool, to perform phasing. Longshot builds on the read-based haplotype phasing algorithm, HapCUT2 [10] and employs a pair-Hidden Markov Model (pair-HMM) to mitigate uncertainties in local alignment. This approach enables the estimation of precise base quality values, which are crucial for genotype likelihood calculations. However, since Longshot is specifically designed to detect and phase single nucleotide variants (SNVs) and only approximately 75% of the reads can be partitioned into two distinct haplotypes, we assume the remaining 25% of reads correspond to highly homogeneous regions. As a result, FocalSV assigns the remaining reads directly to both haplotypes of the nearest phase block.

#### Haplotype-aware local assembly

After reads partitioning, every read within the region BAM file is assigned to a certain phase block and haplotype. FocalSV then performs local assembly based on these phase blocks and haplotypes. For Pacbio Hifi reads, FocalSV adopts Hifiasm (v0.14) [11] for local assembly, while for Pacbio CLR and Nanopore (ONT) reads, Flye (v2.9.1) [12] is utilized. Haplotype-resolved contigs are generated at the end of this procedure.

#### Detection of Candidate indel SVs

The haplotype-resolved contigs are then aligned to the human reference genome using minimap2 [13] and samtools [14] in FocalSV. The example command is as follows:

~~~
minimap2 -a -x asm5 --cs -r2k -t 30 \
       <hg19_ref> \
       <contigs_fasta> \
       | samtools sort >
contigs.bam samtools index contigs.bam
~~~

Notably, the contigs-to-reference BAM file shares similar features with the reads-to-reference BAM file, as both files contain intra- and inter-alignments inferred by CIGAR operations, which indicate potential SVs. However, two key differences exist in terms of coverage and length: the reads-to-reference BAM file typically exhibits much higher coverage, while the contigs are generally much longer than individual reads. To reliably collect SV-related signatures from contig-to-reference BAM file, we adapt the contig-based signature collection methodology from VolcanoSV [15], which refines and optimizes the conventional reads-based SV signature collection methods.

The contig-based signature collection method can be intuitively explained. When an INS is present in the individual’s genome, the contig can be conceptualized as a structure of “reference sequence A + INS sequence + reference sequence B”. If the INS sequence is short, it is typically supported by an intra-alignment, which can be directly extracted from the CIGAR operation. However, longer INS sequences are supported by an inter-alignment (split alignments), where reference sequences A and B align independently to the human reference, while the INS sequence remains unaligned. Similarly, for a DEL in the individual’s genome, the human reference genome can be viewed as having a structure of “sequence A + sequence B + sequence C”, while the corresponding contig can be considered approximately as “sequence A + sequence C”, with sequence B deleted. This deletion can be directly identified from the CIGAR operation. DELs can also be inferred from split alignments, where sequences A and C (adjacent on the contig) are aligned independently to two distant regions on the reference genome.

An INS signature, inferred from a pair of split-alignment events, is collected as follows:

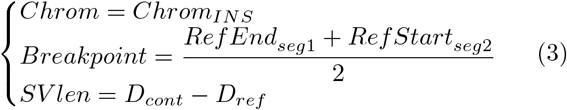

when

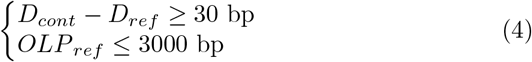

A DEL signature is collected as follows:

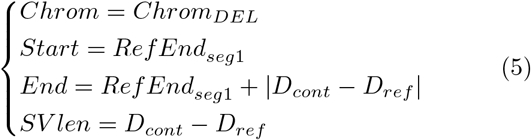

when

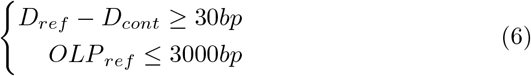

where *Ref Start*_*segi*_ and *Ref End*_*segi*_ denote the start and end coordinates of i-th aligned sequence segment relative to the human reference (*i* ∈ [1, 2]). The segment with the smaller start coordinate on the reference genome is designated as the first aligned segment. *D*_*cont*_ and *D*_*ref*_ represent the distance between two segments relative to the contig and the human reference, respectively. *OLP*_*ref*_ refers to the overlap size between two segments on the human reference.

We then incorporated the clustering pipeline from VolcanoSV [15] to organize the collected signatures by SV type and generate a candidate SV pool. The pipeline involves three steps. Firstly, a chaining clustering algorithm groups signatures from the same haplotype, using SV size similarity and breakpoint distance as clustering criteria. Next, a pairing algorithm matches signatures from different haplotypes for genotyping. Finally, a stringent one-to-K clustering algorithm is applied to reduce the redundancy in the call set.

#### Filtering of indel SVs and refinement of genotypes

To further refine SV calls, FocalSV employs VolcanoSV’s reads-based signature support algorithm [15] to filter out false positive SVs and enhance genotyping precision.

For false positive control, the process begins by gathering read-based SV signatures near the potential SV breakpoints. A similarity score is then calculated between the read-based signature and the candidate SV. Signatures exceeding the similarity threshold are considered supporting signatures. A candidate INS is retained if it has at least one supporting signature, while a candidate DEL must meet the supporting coverage threshold (calculated as the ratio of supporting signatures to local read coverage) to be considered valid. For genotype refinement, a genotype decision tree model [15] is employed to correct the genotype of indel SVs. This model considers five key parameters: contig-based genotype (heterozygous or homozygous), SV size (small or large indel), SV type (insertion or deletion), sequencing technology (Hifi, CLR, or ONT), and a supporting read-based signature ratio (number of supporting read-based signatures/local read depth). With all combinations of the first four parameters, the decision tree consists of 24 leaf nodes. Each leaf node is associated with an empirically determined threshold for the supporting signature ratio. This ratio, ideally reflecting the genotype, is then utilized to predict the final genotype of the SV.

#### Multiple-region mode

FocalSV is also designed to detect SVs across multiple regions. When provided with multiple target regions, FocalSV first retrieves region-specific BAM files, and applies the same steps for each region: reads partitioning, local assembly, and candidate SV detection. This process is parallelized in FocalSV for greater efficiency. Afterward, FocalSV merges all VCF files from each region into a single VCF file. A clustering algorithm is then applied to the merged VCF file to eliminate redundant variants. Finally, the SV filtering and genotype refinement are performed to produce the final VCF file.

### Complex SV detection, recovery, and breakend refinement

We designed specific approaches to recover duplications from insertions and refine the breakends of translocations and inversion.

#### Duplication detection and recovery

For duplication detection, FocalSV integrates information from both contigs and read-based BAM files. The user provides the approximate start and end positions of the target duplication as prior knowledge. Using this input, contigs are generated through a region-based assembly pipeline, the same approach used for large INDEL detection. These contigs are subsequently aligned to a human reference genome to produce a contig-based BAM file. Simultaneously, a read-based BAM file is generated by extracting reads from the user-defined target duplication region along with an extended flanking region (default: 25 kb). In the alignment-based information derived from the contig-based BAM file, duplication (DUP) can be regarded as a special instance of insertion (INS), particularly when the inserted sequence closely resembles a reference segment near the insertion breakpoint. As a result, if two adjacent segments on the contig align to the same region on the reference genome, a duplication can be directly inferred. However, in practice, due to the limitation of existing aligners, only one of the duplicated segments may be correctly aligned to the reference, while the other segment is “skipped”, leading to it being incorrectly labeled as an INS in the BAM file. To address this, we designed a dedicated duplication recovery pipeline from insertion calls. Specifically, FocalSV extracts the alternate alleles from all insertion calls and realigns them to the reference genome. If an alternate allele aligns close to its corresponding insertion breakpoint, it indicates the presence of a duplication (Figure 2A). The duplication’s start and end coordinates are defined by the alignment coordinates of the alternate allele.

This recovery procedure can recover a considerable amount of DUPs missed by the aligner. However, in cases where the contigs are collapsed (i.e. do not contain the duplicated segments) due to misassembly, we designed an additional reads alignment-based approach to recover those DUPs. In this approach, FocalSV identifies events in the target region where a read aligns more than once on the same strand, generating an alignment records pool for each muti-aligned read (MAR). For each MAR, the positional relationship between any two alignment records is evaluated to determine whether they suggest a duplication. A DUP signature is inferred when the pair of read segment alignment records meet the below conditions:

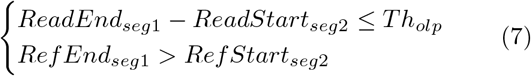

where *ReadStart*_*segi*_, *ReadEnd*_*segi*_, *Ref Start*_*segi*_ and *Ref End*_*segi*_ represent the start and end coordinates of the *i*-th segment relative to read and reference, respectively (Figure 2A). *Th*_*olp*_ denotes the maximum allowed overlap between two segments (500 bp by default). The segments are ordered such that *ReadStart*_*seg*1_ *< ReadStart*_*seg*1_.

Next, FocalSV applies a distance-based clustering algorithm to group DUP signatures. Two DUP signatures are clustered together if their breakpoint shift is less than a specified distance threshold (1000bp by default). The average start and end positions within each cluster are selected as the final breakpoints of the DUP. It is important to note that this reads-based approach is only used for DUPs estimated to be smaller than 5Mb, as split-alignments with gaps larger than 5Mb on the reference are likely misalignments and unreliable. Finally, FocalSV merges the result from both the conitg-based and read-based BAM files to generate the final set of DUP calls.

#### Translocation detection

The format for a typical translocation (TRA) is:

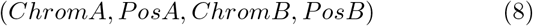

To accurately refine breakends (BND) of TRAs, FocalSV requires prior knowledge of the approximate regions for both BNDs, *PosA* and *PosB*:

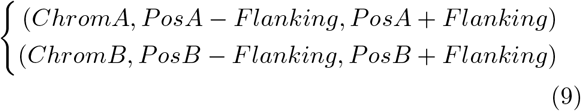

To achieve the detection and refinement, FocalSV identifies TRA signatures from read-based BAM files and calculates the optimal BND. Specifically, TRAs are inferred when two adjacent segments on the read align to different chromosomes. FocalSV first collects reads aligned to both BND regions using equation 9. It then calculates the segment distance between two alignment records of each read. A TRA event is inferred if the two alignment records satisfy the below conditions:

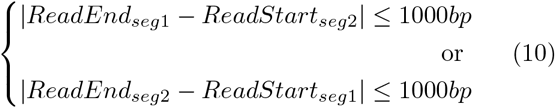

where *ReadStart*_*seg*1_, *ReadEnd*_*seg*1_, *ReadStart*_*seg*2_, and *ReadEnd*_*seg*2_ represent the start and end positions of each aligned segment on the read, respectively. A TRA signature is represented as follows:

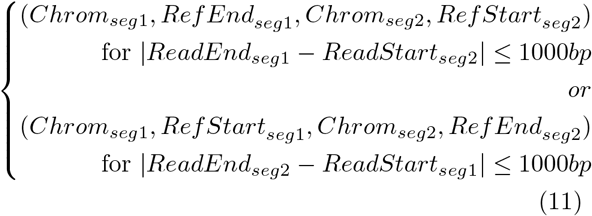

where *Ref Start*_*seg*1_, *Ref End*_*seg*1_, *Ref Start*_*seg*2_, and *Ref End*_*seg*2_ denote the start and end positions of each aligned segment on the reference genome, respectively (Figure 2B). A clustering algorithm is then employed to merge TRA signatures. Any two TRA signatures are grouped into a single cluster if they meet the following criteria:

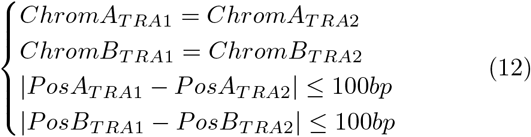

The average coordinate for each BND is selected as the cluster center, serving as the final TRA call.

#### Inversion detection

The format for a typical inversion (INV) is:

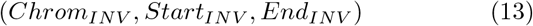

Similar as TRA, the target region for INV is defined as follows:

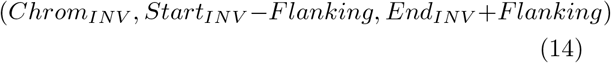

FocalSV then identifies INV signatures from the read-based BAM file. Specifically, INV events are identified when two adjacent segments of a read align to two distant locations on the reference in reverse orientations. The criteria for inferring an INV and its corresponding signature are outlined below:

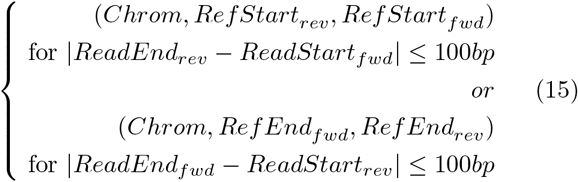

where *ReadStart*_*fwd*_, *ReadEnd*_*fwd*_, *ReadStart*_*rev*_, and *ReadEnd*_*rev*_ denote the start and end positions of the forward- and reverse-aligned segment on the read, while *Ref Start*_*fwd*_, *Ref End*_*fwd*_, *Ref Start*_*rev*_, and *Ref End*_*rev*_ represent the start and end positions on the reference (Figure 2C). A clustering algorithm is then applied to merge INV signatures. Two INV signatures are grouped into a single cluster if they meet the following criteria:

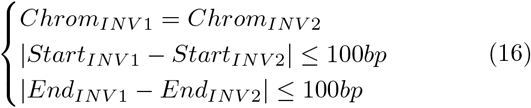

The average position for each BND is selected as the cluster center, serving as the final INV call.

### Somatic SV discovery in cancer genome

To assess the effectiveness of various SV calling methods in identifying somatic mutations, we utilized Pacbio and ONT Tumor-Normal paired libraries of sample HCC1395. We also incorporated the high-confidence HCC1395 somatic SV call set as a gold standard, following the methodology outlined by Talsania et al [16]. Initially, SV calls from the paired libraries were generated by each method. Subsequently, the SV filtering and merging tool SURVIVOR [17] was employed to extract somatic mutations. The resulting somatic SV call sets were then compared to the high-confidence somatic SV call set to assess performance. The pipeline for employing SURVIVOR to identify somatic mutations is outlined as follows. We first segmented VCF files according to SV type (translocation and non-translocation) and size windows (for non-translocations). The size windows recommended by Talsania et al. are as follows: 50-100bp, 101-500bp, 501-1000bp, 1001-30000bp, *>*30000bp. We utilized the following SURVIVOR command to split each VCf file:

~~~
SURVIVOR filter <input_vcf> \
NA <min_sv_size> <max_sv_size> 0 -1 \
<out_vcf>
~~~

Next, somatic mutations were extracted by merging the tumor and normal SV call sets. Each VCF file of a specific size window employed the lower bound of its size window as the maximum threshold of break-point distance when merging SVs. For instance, in a 50-100bp VCF file, the threshold for breakpoint distance was set to 50bp. This strategy ensures consistency and accuracy in the merging process across different size windows.

~~~
SURVIVOR merge <VCFlist> \
<dist_thresh> 1 1 0 0 <min_sv_size> \
<merged_vcf>
~~~

The *<*VCFlist*>* denotes the VCF file path for the normal and tumor SV call sets. Following the merging procedure, SVs supported exclusively by the tumor call set, without corresponding support in the normal call set, were filtered out as somatic SVs. This filtering procedure was executed on non-TRA VCF files across various size windows. Subsequently, the five resulting filtered VCF files for non-TRA plus the TRA VCF file were concatenated into a single VCF file representing the final somatic mutation call set.

In the final step, the evaluation of somatic mutations against the high-confidence gold standard call set was conducted using distinct criteria. Specifically, for TRAs, assessment was carried out solely at the breakend level. The breakend shift (r) between TRAs from the call set and the gold standard set was calculated. If the value of r is less than or equal to 1kb, the TRA is classified as true positive (TP); otherwise, it is labeled as false positive (FP). For non-TRA SVs, both the breakend shift (r) and size similarity (P) are calculated.

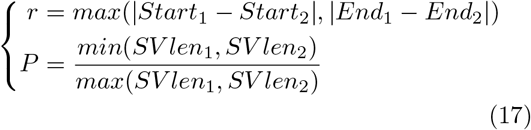

For non-TRA SVs, if the value of r is less than or equal to 500bp and P is greater than or equal to 0.5, the non-TRA SV is classified as true positive (TP); otherwise, it is categorized as false positive (FP). This stringent assessment approach ensures the robustness and accuracy of the evaluation process, effectively distinguishing true somatic mutations from potential artifacts or false positives.

## Results

FocalSV leverages an innovative, region-based, haplotype-aware assembly strategy, providing a thorough pipeline for structural variant (SV) detection, filtering, and refinement by integrating contig-level assembly data and read-level information (Figure 1 2). This sophisticated approach enables FocalSV to generate accurate and comprehensive SV call sets. To demonstrate the superior performance of FocalSV, we benchmarked it against four state-of-the-art assembly-based and five alignment-based SV detection tools across multiple long-read sequencing datasets. The assembly-based tools evaluated include FocalSV (v1.0.0), PAV (freeze2) [18], SVIM-asm (v1.0.2) [19], and Dipcall (v0.3) [20], while the alignment-based tools comprise CuteSV (v1.0.11) [21], SVIM (v1.4.2) [22], pbsv (v2.6.2) [23], Sniffles2 (v2.0.6) [24], and SKSV (v1.0.2) [25].

**Figure 1.**
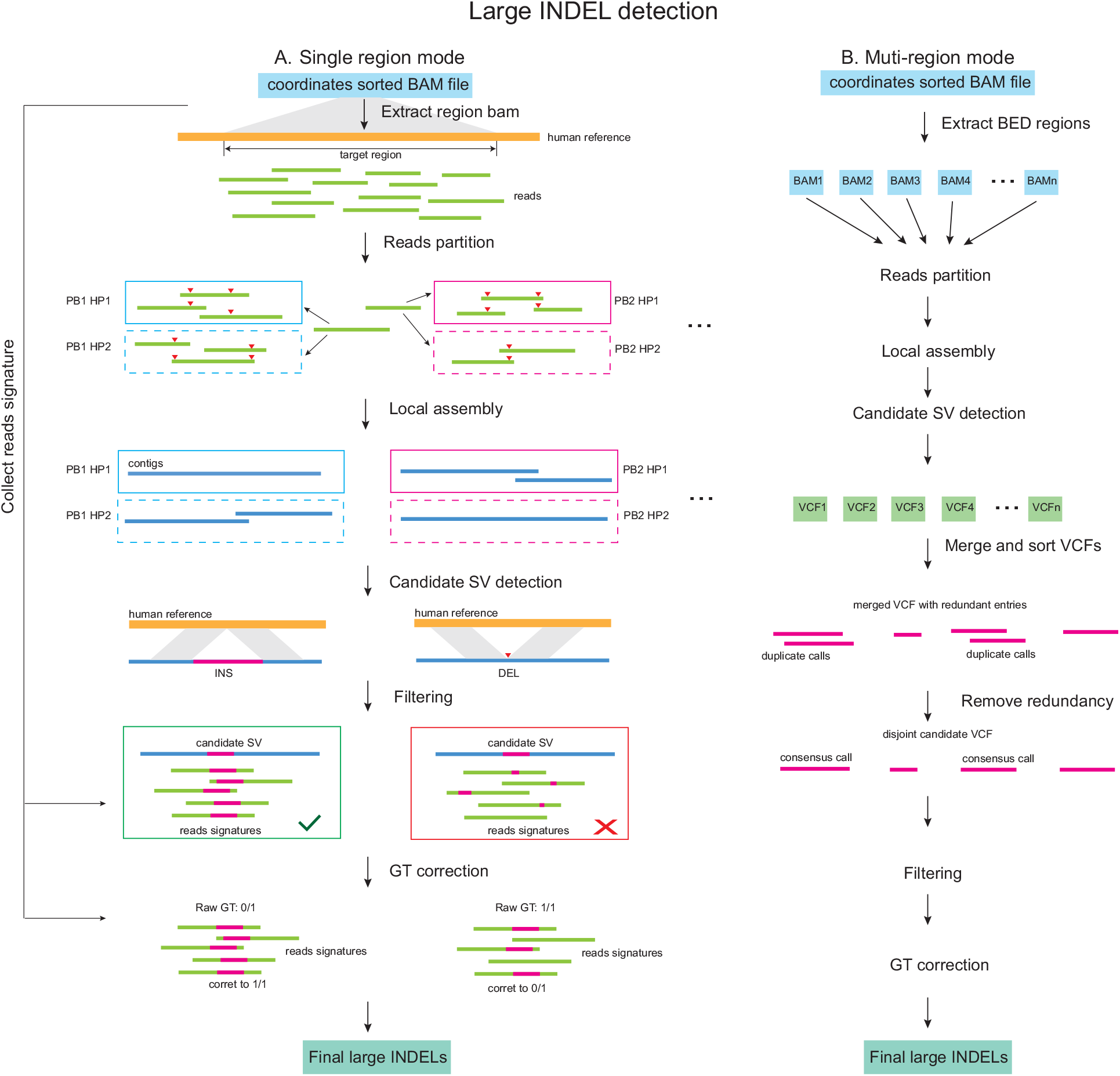
Schematic diagram of the FocalSV large indel detection pipeline. The workflow for large indel detection includes two modes: **(a)** single region mode and **(b)** multi-region mode. **(a) Single region mode:** the input data include a high-quality reference genome and a BAM file containing aligned long reads. The reads extraction module isolates long reads aligned to the region of interest. The haplotyping module partitions these reads into distinct parental haplotypes. The local assembly module uses the phased reads to perform independent de novo local assemblies. Finally, the variant calling module identifies indel structural variants (SVs) by comparing the assembled contigs to the reference genome, followed by filtering and genotype (GT) correction in postprocessing steps. **(b) Multi-region mode:** the input data include a high-quality reference genome and a BAM file containing aligned long reads. FocalSV retrieves region-specific BAM files and processes each region independently through reads partitioning, local assembly, and SV detection. The VCF files from all regions are merged into a single file, and redundant variants are removed using a clustering algorithm. SV filtering and genotype refinement are then applied to produce the final VCF file.

**Figure 2.**
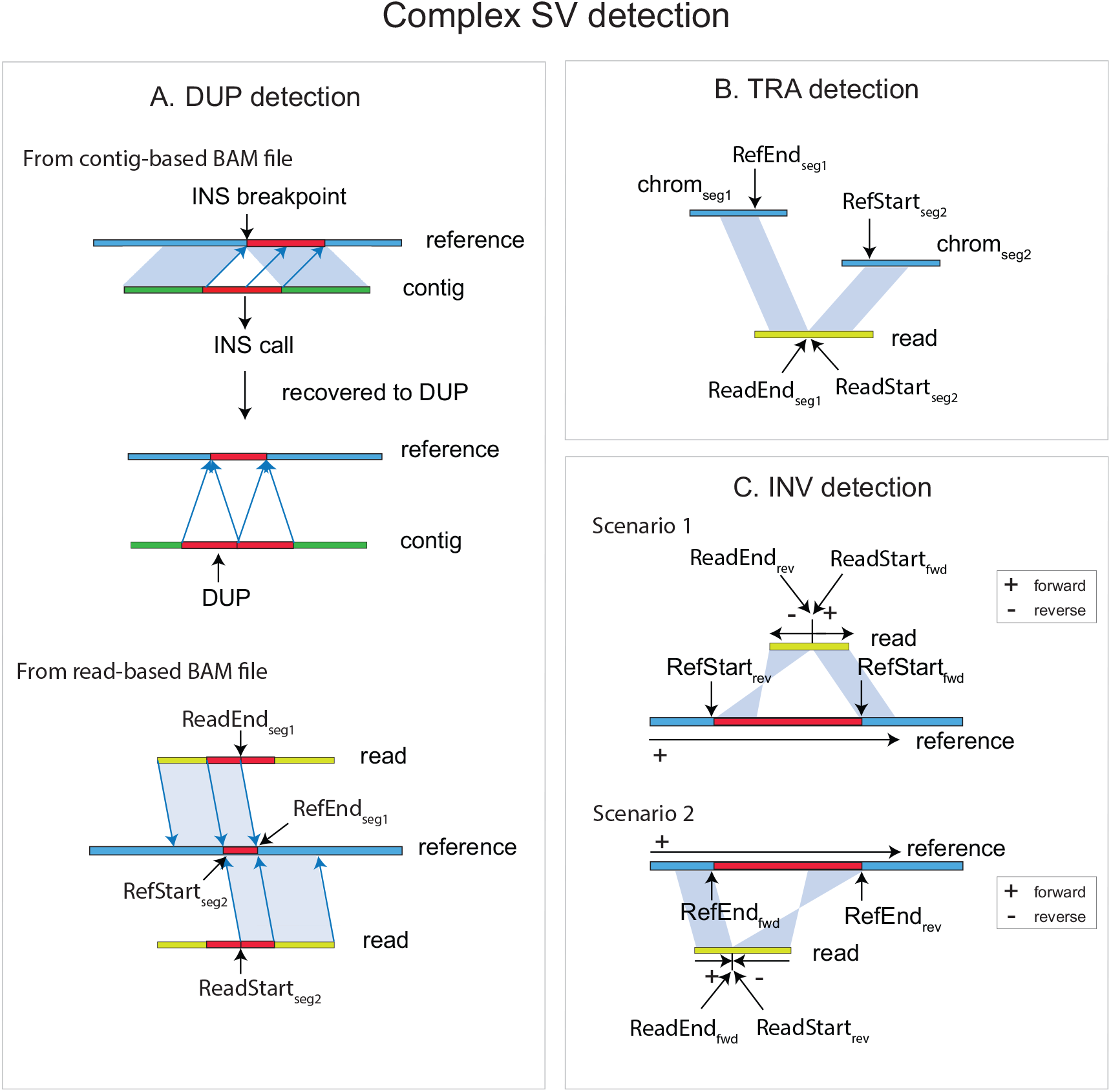
Schematic diagram of complex SV signatures detected by FocalSV. **(a) Duplication:** duplications can be identified using both contig-based and read-based BAM files. In a contig-based BAM file, a duplication is detected when an insertion call shows the alternate allele mapped to the surrounding sequence of the insertion breakpoint. In a read-based BAM file, a duplication signature is identified when two adjacent segments of a read align to overlapping regions on the reference genome in the same orientation. **(b) Translocation:** translocations are inferred when two adjacent segments of a read align to different chromosomes. **(c) Inversion:** inversions are detected when two adjacent segments of a read align in opposite orientations. Additional details on detecting complex SVs can be found in the Methods section.

The datasets include nine libraries derived from the well-characterized HG002 sample, sequenced with PacBio HiFi, CLR, and ONT technologies, as well as two paired tumor-normal datasets (CLR and ONT) from the HCC1395 cancer cell line. Specifically, the HG002 datasets encompass three HiFi libraries (Hifi L1, Hifi L2, and Hifi L3) with coverages ranging from 30× to 56×, three CLR libraries (CLR L1, CLR L2, and CLR L3) with coverages between 29× and 89×, and three ONT libraries (ONT L1, ONT L2, and ONT L3) with coverages between 47× and 57×. Further details on the tools and datasets are provided in Table 1.

**Table 1.**
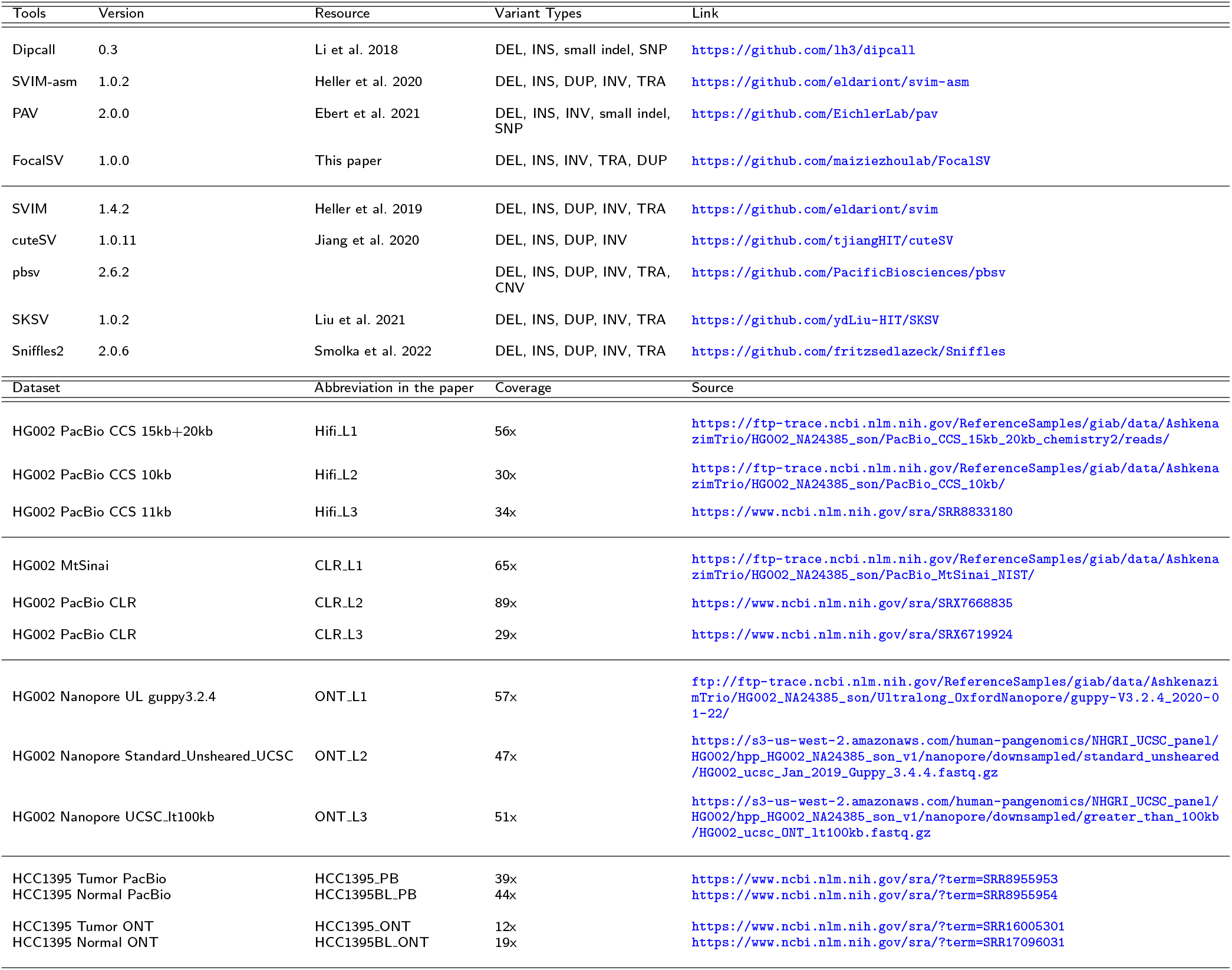
Resource for different tools and long-read datasets. Top panel: The SV calling tools used in this paper. Tool version number, cited article, variant types called by each tool, and tool links are shown in the table. Bottom panel: The long-read datasets used in this paper. The abbreviation, coverage, and source link for each dataset are shown in the table. The abbreviation is used to refer to each dataset in the main text and supplementary information.

We assessed the performance of all tools based on key metrics, including breakpoint identification and SV sequence accuracy. FocalSV’s pipeline, as illustrated in Figure 1, demonstrates how its region-based assemblies generate high-quality contigs that facilitate precise SV detection. Through a rigorous filtering process, FocalSV ensures that the final SV call sets are both exhaustive and accurate, reflecting the tool’s ability to capture a broad range of SVs. The complete methodology and implementation details are described in the Methods section.

### Evaluation of SV calls in FocalSV and comparisons with existing SV callers

To assess the performance of insertion and deletion SV detection, we evaluated four assembly-based tools (FocalSV, PAV, SVIM-asm, and Dipcall) and five alignment-based tools (CuteSV, SVIM, pbsv, Sniffles2, and SKSV) using nine long-read sequencing libraries of the HG002 sample. We benchmarked the results against the Genome in a Bottle (GIAB) SV gold standard [26] using Truvari (v4.0.0) [27], a widely adopted structural variant evaluation tool. Truvari compares SV calls from any tool’s call set against a gold standard call set, both provided in Variant Call Format (VCF) files, by analyzing key metrics: reference distance, reciprocal overlap, size similarity, and sequence similarity.

In this study, we applied a moderate-tolerance parameter set in Truvari for SV comparisons, with the following settings: p=0.5, P=0.5, r=500, and O=0.01. Specifically, the parameter p (pctstim), which ranges from 0 to 1.0, controls the minimum required sequence similarity for two SVs to be considered identical. P (pctsize), also ranging from 0 to 1.0, defines the minimum allowable allele size similarity. O (pctovl), ranging from 0 to 1.0, establishes the minimum reciprocal overlap ratio, a key measure for comparing deletions and evaluating their breakpoint alignment. Lastly, r (refdist), which can range from 0 to 1000bp, sets the maximum allowable difference between the reference positions of two SVs, helping assess breakpoint shifts in insertion events.

We first evaluated the average performance across different PacBio Hifi, CLR, and ONT datasets (Table 2-4). In the HiFi datasets (Table 2), FocalSV demonstrated the highest average F1 scores for both deletions (93.99%) and insertions (91.93%), while also achieving the best genotyping accuracy for deletions (99.03%) and the second-best accuracy for insertions (98.30%). In the CLR datasets (Table 3), FocalSV outperformed the other tools with the highest average F1 scores for deletions (92.95%) and insertions (90.71%), although it ranked third for genotyping accuracy in deletions (98.22%) and second for insertions (96.34%). In the ONT datasets (Table 4), FocalSV once again achieved the highest average F1 score for deletions (92.94%), ranked second for insertions (89.69%), and secured the second-best genotyping accuracy for both deletions (99.06%) and insertions (98.27%). These results highlight the robustness of FocalSV across different sequencing platforms, particularly for deletion detection, where it consistently excels.

When examining each dataset individually (Table 2-4 and Table S1-S3), FocalSV outperformed all other tools, achieving the highest F1 scores for both deletions and insertions across all Hifi libraries, for insertions across all CLR libraries, and deletions across all ONT libraries. In the context of three Hifi datasets (Table 2 and Table S1), FocalSV emerged as the top overall performer. For deletions, FocalSV outperformed all other tools across all libraries, with its F1 score and genotype concordance exceeding the second-ranked tool by an average of 0.36% and 0.27%, respectively. For deletion recall, on Hifi L1, FocalSV outperformed the second-ranked tool by 1.46%. On Hifi L2 and Hifi L3, FocalSV achieved the second-highest recall, trailing the top recall by an average of 0.27%. In terms of deletion precision, FocalSV outperformed the second-ranked tools on Hifi L1 by 0.11%. On Hifi L2 and Hifi L3, it ranked third in precision, with an average of 1.37% less than the highest precision. For insertions, FocalSV outperformed all other tools across all libraries, with F1 score and recall exceeding the second-ranked tools by an average of 1.80% and 0.63%, respectively. In terms of insertion precision, FocalSV outperformed the second-ranked tool on Hifi_L1 by 0.8%. On Hifi_L2 and Hifi_L3, FocalSV achieved the third- and second-highest precision, with an average of 2% less than the top precision.

**Table 2.** Large deletions (DELs) and insertions (INSs) (≥50bp) calling performance across three Hifi datasets. The table presents recall, precision, F1 score (in %), and genotype accuracy (measured by Genotype Concordance) for all benchmarked SV callers. The highest scores for recall, precision, F1, and genotype accuracy are highlighted in bold. Additionally, the overall average performance for each SV type and library is summarized at the bottom of the table. Benchmarking was conducted using Truvari with the following parameter settings: p = 0.5, P = 0.5, r = 500, and O = 0.01.

|  | Library | Metric | FocalSV | PAV | SVIM-asm | Dipcall | cuteSV | SVIM | PBSV | Sniffles2 | SKSV |
| --- | --- | --- | --- | --- | --- | --- | --- | --- | --- | --- | --- |
| DEL | Hifi_L1 | Precision | **93.14%** | 92.28% | 91.70% | 90.33% | 88.20% | 87.33% | 93.03% | 83.75% | 85.29% |
|  |  | Recall | **95.36%** | 93.22% | 93.90% | 92.61% | 88.46% | 88.78% | 93.44% | 84.65% | 85.25% |
|  |  | F1 | **94.24%** | 92.75% | 92.79% | 91.46% | 88.33% | 88.05% | 93.24% | 84.20% | 85.27% |
|  |  | Genotype Concordance | **99.10%** | 97.11% | 97.85% | 96.75% | 99.07% | 98.71% | 97.71% | 96.56% | 97.98% |
|  | Hifi_L2 | Precision | 92.65% | 92.04% | 92.01% | 90.11% | **93.83%** | 82.90% | 92.21% | 92.67% | 85.81% |
|  |  | Recall | 95.00% | 94.12% | 94.56% | 92.76% | 87.12% | **95.40%** | 93.78% | 94.85% | 78.91% |
|  |  | F1 | **93.81%** | 93.07% | 93.27% | 91.42% | 90.35% | 88.70% | 92.99% | 93.74% | 82.22% |
|  |  | Genotype Concordance | **99.00%** | 97.26% | 98.61% | 96.86% | 97.46% | 97.90% | 98.19% | 98.31% | 97.48% |
|  | Hifi_L3 | Precision | 92.67% | 91.71% | 91.70% | 89.94% | **94.23%** | 75.30% | 92.49% | 92.81% | 85.88% |
|  |  | Recall | 95.21% | 93.49% | 94.44% | 92.76% | 90.43% | **95.34%** | 93.93% | 95.02% | 81.44% |
|  |  | F1 | **93.92%** | 92.59% | 93.05% | 91.33% | 92.29% | 84.14% | 93.20% | 93.90% | 83.60% |
|  |  | Genotype Concordance | **98.98%** | 96.62% | 98.59% | 96.83% | 98.04% | 98.34% | 98.60% | 98.47% | 98.06% |
|  | Overall | Precision | **92.82%** | 92.01% | 91.80% | 90.13% | 92.09% | 81.84% | 92.58% | 89.74% | 85.66% |
|  |  | Recall | **95.19%** | 93.61% | 94.30% | 92.71% | 88.67% | 93.17% | 93.72% | 91.51% | 81.87% |
|  |  | F1 | **93.99%** | 92.80% | 93.04% | 91.40% | 90.32% | 86.96% | 93.14% | 90.61% | 83.70% |
|  |  | Genotype Concordance | **99.03%** | 97.00% | 98.35% | 96.81% | 98.19% | 98.32% | 98.17% | 97.78% | 97.84% |
| INS | Hifi_L1 | Precision | **90.80%** | 86.30% | 86.50% | 72.80% | 87.50% | 85.10% | 90.00% | 79.00% | 86.10% |
|  |  | Recall | **93.90%** | 93.10% | 93.10% | 91.40% | 82.30% | 69.60% | 77.40% | 76.80% | 86.60% |
|  |  | F1 | **92.40%** | 89.60% | 89.70% | 81.00% | 84.90% | 76.60% | 83.20% | 77.90% | 86.40% |
|  |  | Genotype Concordance | 98.10% | 95.60% | 97.80% | 77.80% | **99.10%** | 90.00% | 87.70% | 89.30% | 98.30% |
|  | Hifi_L2 | Precision | 89.80% | 86.40% | 86.40% | 73.10% | **91.80%** | 61.90% | 89.90% | 88.30% | 86.20% |
|  |  | Recall | **93.80%** | 93.30% | 92.90% | 91.60% | 82.40% | 91.60% | 76.70% | 91.80% | 61.70% |
|  |  | F1 | **91.70%** | 89.70% | 89.50% | 81.30% | 86.80% | 73.90% | 82.80% | 90.00% | 71.90% |
|  |  | Genotype Concordance | **98.30%** | 96.10% | 97.70% | 78.50% | 97.60% | 94.00% | 95.40% | 97.00% | 97.70% |
|  | Hifi_L3 | Precision | 89.60% | 86.30% | 86.40% | 73.00% | **91.60%** | 45.00% | 89.00% | 88.60% | 87.10% |
|  |  | Recall | **93.80%** | 93.00% | 93.20% | 91.80% | 86.00% | 92.10% | 76.70% | 92.80% | 71.00% |
|  |  | F1 | **91.70%** | 89.50% | 89.60% | 81.30% | 88.70% | 60.50% | 82.40% | 90.70% | 78.20% |
|  |  | Genotype Concordance | 98.50% | 94.90% | 98.00% | 78.30% | 98.40% | 94.30% | 96.20% | 97.10% | **98.60%** |
|  | Overall | Precision | 90.07% | 86.33% | 86.43% | 72.97% | **90.30%** | 64.00% | 89.63% | 85.30% | 86.47% |
|  |  | Recall | **93.83%** | 93.13% | 93.07% | 91.60% | 83.57% | 84.43% | 76.93% | 87.13% | 73.10% |
|  |  | F1 | **91.93%** | 89.60% | 89.60% | 81.20% | 86.80% | 70.33% | 82.80% | 86.20% | 78.83% |
|  |  | Genotype Concordance | 98.30% | 95.53% | 97.83% | 78.20% | **98.37%** | 92.77% | 93.10% | 94.47% | 98.20% |

In the three CLR datasets (Table 3 and Table S2), FocalSV emerged as the top-performing tool in terms of F1 score and recall across nearly all libraries, demonstrating clear advantages. For deletions, FocalSV outperformed the second-ranked tool in F1 score on CLR L1 and CLR L3 by an average of 0.30%. On CLR L2, FocalSV exhibited the second-highest F1 score, trailing the top by 0.19%. For deletion recall, FocalSV outperformed the second-ranked tools across all three libraries by an average of 1.48%. However, in terms of deletion precision, FocalSV’s performance was slightly lower, ranking 5th, 4th, and 4th on CLR L1, CLR L2, and CLR L3, respectively, with an average of 1.74% less than the top precision. For deletion genotype concordance, FocalSV outperformed the second-ranked tool on CLR L1 by 0.08%. On CLR L2 and CLR L3, FocalSV achieved the second- and third-highest genotype concordance, trailing the top by an average of 1.07%. For insertions, FocalSV excelled across all libraries, with its F1 surpassing the second-ranked tools by an average of 3.45%. For insertion recall, FocalSV achieved the second-ranked tools on CLR L1 and CLR L2 by an average of 0.93%. On CLR L3, FocalSV exhibited the second-highest recall, trailing the top by 0.27%. In terms of insertion precision, FocalSV outperformed the second-ranked tool on CLR L2 by 1.88%. On CLR L1 and CLR L3, it ranked third and second, respectively, with an average of 3.04% less than the top precision. For insertion genotype concordance, FocalSV exhibited the second-highest genotype concordance across CLR L1, CLR L2, and CLR L3, with an average of 1.40% less than the top genotype concordance.

**Table 3.** Large deletions (DELs) and insertions (INSs) (≥50bp) calling performance across three CLR datasets. The table presents recall, precision, F1 score (in %), and genotype accuracy (measured by Genotype Concordance) for all benchmarked SV callers. The highest scores for recall, precision, F1, and genotype accuracy are highlighted in bold. Additionally, the overall average performance for each SV type and library is summarized at the bottom of the table. Benchmarking was conducted using Truvari with the following parameter settings: p = 0.5, P = 0.5, r = 500, and O = 0.01.

|  |  |  | FocalSV | PAV | SVIM-asm | Dipcall | cuteSV | SVIM | PBSV | Sniffles2 |
| --- | --- | --- | --- | --- | --- | --- | --- | --- | --- | --- |
|  | Library | Metric |  |  |  |  |  |  |  |  |
| DEL | CLR.L1 | Precision | 91.76% | 81.51% | 88.51% | 82.40% | 92.04% | 91.80% | **93.99%** | 92.49% |
|  |  | Recall | **94.17%** | 86.88% | 86.10% | 21.04% | 89.38% | 91.70% | 91.89% | 91.81% |
|  |  | F1 | **92.95%** | 84.11% | 87.29% | 33.52% | 90.69% | 91.70% | 92.92% | 92.15% |
|  |  | Genotype Concordance | **98.76%** | 96.00% | 95.23% | 85.45% | 98.64% | 97.90% | 96.64% | 98.68% |
|  | CLR.L2 | Precision | 92.03% | 91.21% | 92.01% | 88.81% | 91.43% | **93.40%** | 93.29% | 92.93% |
|  |  | Recall | **94.83%** | 89.02% | 91.72% | 86.95% | 91.98% | 93.90% | 92.95% | 92.23% |
|  |  | F1 | 93.41% | 90.10% | 91.86% | 87.87% | 91.70% | **93.60%** | 93.12% | 92.57% |
|  |  | Genotype Concordance | 98.72% | 98.39% | 98.46% | 94.36% | **99.29%** | 98.60% | 97.80% | 97.37% |
|  | CLR.L3 | Precision | 92.45% | 88.32% | 90.60% | 83.06% | 92.22% | 93.00% | 92.56% | **94.08%** |
|  |  | Recall | **92.54%** | 84.35% | 84.28% | 46.33% | 80.10% | 79.30% | 91.30% | 78.33% |
|  |  | F1 | **92.50%** | 86.29% | 87.33% | 59.48% | 85.74% | 85.60% | 91.93% | 85.48% |
|  |  | Genotype Concordance | 97.24% | 95.65% | 96.17% | 84.27% | 97.76% | **98.80%** | 96.78% | 96.00% |
|  | Overall | Precision | 92.08% | 87.01% | 90.37% | 84.76% | 91.90% | 92.73% | **93.28%** | 93.17% |
|  |  | Recall | **93.85%** | 86.75% | 87.37% | 51.44% | 87.15% | 88.30% | 92.05% | 87.46% |
|  |  | F1 | **92.95%** | 86.83% | 88.83% | 60.29% | 89.38% | 90.30% | 92.66% | 90.07% |
|  |  | Genotype Concordance | 98.24% | 96.68% | 96.62% | 88.03% | **98.56%** | 98.43% | 97.07% | 97.35% |
| INS | CLR.L1 | Precision | 89.15% | 72.05% | 80.21% | 69.61% | 86.78% | 92.73% | **93.15%** | 88.02% |
|  |  | Recall | **92.84%** | 91.18% | 89.95% | 20.77% | 86.12% | 71.20% | 79.78% | 86.65% |
|  |  | F1 | **90.96%** | 80.49% | 84.80% | 32.00% | 86.45% | 80.55% | 85.94% | 87.33% |
|  |  | Genotype Concordance | 96.94% | 91.75% | 92.00% | 68.00% | **98.04%** | 61.70% | 87.56% | 96.72% |
|  | CLR.L2 | Precision | **87.98%** | 84.15% | 84.62% | 66.76% | 83.15% | 62.85% | 86.10% | 81.14% |
|  |  | Recall | **93.52%** | 91.76% | 93.32% | 88.13% | 91.88% | 27.65% | 77.90% | 80.89% |
|  |  | F1 | **90.67%** | 87.79% | 88.75% | 75.97% | 87.30% | 38.40% | 81.80% | 81.02% |
|  |  | Genotype Concordance | 97.02% | 95.87% | 96.77% | 67.60% | **98.97%** | 44.11% | 89.52% | 94.97% |
|  | CLR.L3 | Precision | 89.59% | 78.91% | 81.51% | 60.40% | 88.16% | **91.67%** | 87.57% | 89.51% |
|  |  | Recall | 91.44% | **91.71%** | 90.40% | 47.28% | 79.83% | 75.48% | 79.93% | 70.42% |
|  |  | F1 | **90.51%** | 84.83% | 85.72% | 53.04% | 83.79% | 82.79% | 83.58% | 78.83% |
|  |  | Genotype Concordance | 95.07% | 89.24% | 89.86% | 54.27% | **96.23%** | 87.68% | 88.65% | 92.63% |
|  | Overall | Precision | 88.91% | 78.37% | 82.11% | 65.59% | 86.03% | 82.42% | **88.94%** | 86.22% |
|  |  | Recall | **92.60%** | 91.55% | 91.22% | 52.06% | 85.94% | 58.11% | 79.20% | 79.32% |
|  |  | F1 | **90.71%** | 84.37% | 86.42% | 53.67% | 85.85% | 67.25% | 83.77% | 82.39% |
|  |  | Genotype Concordance | 96.34% | 92.29% | 92.88% | 63.29% | **97.75%** | 64.50% | 88.58% | 94.77% |

In the three ONT datasets (Table 4 and Table S3), FocalSV maintained substantial leads. It outperformed the second-ranked tool across ONT L1, ONT L2, and ONT L3 by an average of 0.58% and 0.84% in deletion F1 score and precision, respectively. For deletion recall, FocalSV outperformed the second-ranked tools on ONT L1 and ONT L2 by an average of 0.58%. On ONT L3, FocalSV achieved the second-highest recall, trailing the top by 0.10%. For deletion genotype concordance, FocalSV outperformed the second-ranked tool on ONT L2 by 0.09%. On ONT L1 and ONT L3, FocalSV achieved the third- and second-highest genotype concordance, with an average of 0.22% less than the top genotype concordance. For insertion F1, FocalSV ranked third, second, and second on ONT L1, ONT L2, and ONT L3, with an average of 0.91% less than the top F1. In terms of insertion recall, FocalSV outperformed the second-ranked tool on ONT L2 and ONT L3 by an average of 0.51%. On ONT L1, FocalSV achieved the second-highest recall, trailing the top recall by 0.49%. In terms of insertion precision, FocalSV ranked third on ONT L1, ONT L2, and ONT L3, averaging 3.66% below the top precision. For insertion genotype concordance, FocalSV exhibited the third on ONT L1, and second on both ONT L2 and ONT L3, with an average difference of 0.51% below the highest genotype concordance.

**Table 4.** Large deletions (DELs) and insertions (INSs) (≥50bp) calling performance across three ONT datasets. The table presents recall, precision, F1 score (in %), and genotype accuracy (measured by Genotype Concordance) for all benchmarked SV callers. The highest scores for recall, precision, F1, and genotype accuracy are highlighted in bold. Additionally, the overall average performance for each SV type and library is summarized at the bottom of the table. Benchmarking was conducted using Truvari with the following parameter settings: p = 0.5, P = 0.5, r = 500, and O = 0.01.

|  |  |  | FocalSV | PAV | SVIM-asm | Dipcall | cuteSV | SVIM | Sniffles2 |
| --- | --- | --- | --- | --- | --- | --- | --- | --- | --- |
|  | Library | Metric |  |  |  |  |  |  |  |
| DEL | ONT_L1 | Precision | **91.89%** | 91.10% | 90.86% | 86.48% | 91.32% | 90.88% | 91.60% |
|  |  | Recall | **93.90%** | 92.78% | **93.90%** | 90.72% | 92.78% | 93.00% | 92.44% |
|  |  | F1 | **92.89%** | 91.94% | 92.35% | 88.55% | 92.05% | 91.93% | 92.02% |
|  |  | Genotype Concordance | 98.94% | 98.74% | 98.78% | 92.64% | 99.08% | 98.30% | **99.19%** |
|  | ONT_L2 | Precision | **92.03%** | 90.88% | 91.05% | 84.66% | 85.56% | 86.35% | 79.67% |
|  |  | Recall | **93.97%** | 89.04% | 93.71% | 88.61% | 92.40% | 92.54% | 92.35% |
|  |  | F1 | **92.99%** | 89.95% | 92.36% | 86.59% | 88.84% | 89.34% | 85.54% |
|  |  | Genotype Concordance | **99.12%** | 98.72% | 98.73% | 90.18% | 98.87% | 98.03% | 99.03% |
|  | ONT_L3 | Precision | **92.27%** | 90.96% | 91.03% | 85.39% | 88.84% | 88.84% | 88.06% |
|  |  | Recall | 93.61% | 90.89% | **93.71%** | 89.14% | 92.42% | 92.81% | 92.42% |
|  |  | F1 | **92.93%** | 90.92% | 92.35% | 87.22% | 90.59% | 90.78% | 90.18% |
|  |  | Genotype Concordance | 99.12% | 98.74% | 98.81% | 91.06% | 99.11% | 98.46% | **99.32%** |
|  | Overall | Precision | **92.06%** | 90.98% | 90.98% | 85.51% | 88.57% | 88.69% | 86.44% |
|  |  | Recall | **93.83%** | 90.90% | 93.77% | 89.49% | 92.53% | 92.78% | 92.40% |
|  |  | F1 | **92.94%** | 90.94% | 92.35% | 87.45% | 90.49% | 90.68% | 89.25% |
|  |  | Genotype Concordance | 99.06% | 98.73% | 98.77% | 91.29% | 99.02% | 98.26% | **99.18%** |
| INS | ONT_L1 | Precision | 86.38% | 86.03% | 85.28% | 65.47% | **90.32%** | 82.60% | 89.00% |
|  |  | Recall | 92.84% | 92.71% | **93.33%** | 89.45% | 91.55% | 88.45% | 91.14% |
|  |  | F1 | 89.50% | 89.25% | 89.12% | 75.60% | **90.93%** | 85.42% | 90.06% |
|  |  | Genotype Concordance | 98.14% | 97.22% | 98.19% | 63.74% | **98.88%** | 85.70% | 95.95% |
|  | ONT_L2 | Precision | 86.24% | 84.72% | 85.06% | 63.83% | **89.86%** | 81.65% | 88.40% |
|  |  | Recall | **93.60%** | 89.24% | 92.90% | 86.80% | 90.57% | 86.86% | 90.19% |
|  |  | F1 | 89.77% | 86.92% | 88.80% | 73.56% | **90.21%** | 84.17% | 89.29% |
|  |  | Genotype Concordance | 98.30% | 97.18% | 97.92% | 59.25% | **98.49%** | 82.52% | 95.28% |
|  | ONT_L3 | Precision | 86.81% | 84.78% | 84.48% | 64.42% | **90.24%** | 81.55% | 88.65% |
|  |  | Recall | **92.99%** | 90.72% | 92.67% | 86.99% | 91.06% | 88.87% | 90.70% |
|  |  | F1 | 89.80% | 87.65% | 88.39% | 74.03% | **90.65%** | 85.05% | 89.67% |
|  |  | Genotype Concordance | 98.37% | 97.10% | 98.10% | 60.58% | **98.96%** | 86.96% | 95.64% |
|  | Overall | Precision | 86.48% | 85.18% | 84.94% | 64.57% | **90.14%** | 81.93% | 88.68% |
|  |  | Recall | **93.14%** | 90.89% | 92.97% | 87.75% | 91.06% | 88.06% | 90.68% |
|  |  | F1 | 89.69% | 87.94% | 88.77% | 74.40% | **90.60%** | 84.88% | 89.67% |
|  |  | Genotype Concordance | 98.27% | 97.17% | 98.07% | 61.19% | **98.78%** | 85.06% | 95.62% |

**Table 5.**
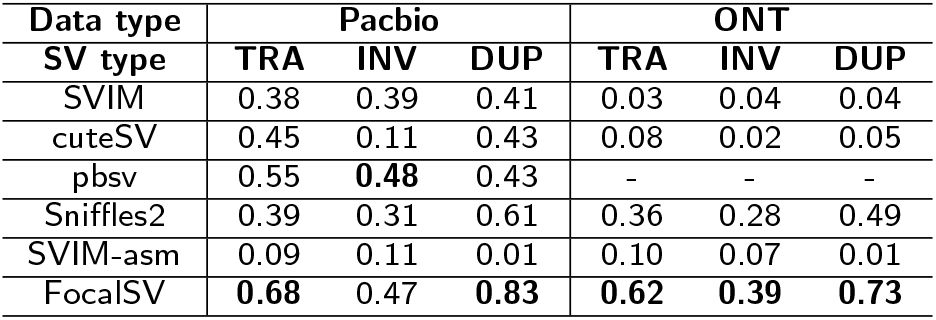
Somatic SV detection accuracy for paired normal-tumor cancer data from Pacbio and ONT sequencing. The table shows recall values for three different types of complex SVs, evaluated across benchmarked tools: translocations (TAR), inversions (INV), and duplication (DUP). Recall values were calculated by assuming SVs from the high-confidence set were the gold standard.

In summary, FocalSV consistently outperformed other tools across nine datasets (Hifi, CLR, and ONT) in terms of F1 score and recall, demonstrating its strength for both deletions and insertions. While it occasionally ranked second or third in precision and genotype concordance, these minor variations did not detract from its overall performance. FocalSV proved to be a reliable and high-performing tool.

### FocalSV demonstrates excellent resilience to different SV size ranges

So far, we evaluated SVs by averaging across all size ranges. However, we found that the tools we assessed showed varying levels of accuracy in detecting SVs of different sizes. To demonstrate the effect of SV size and compare the resilience of each tool, we plotted the F1 score against the range of SV sizes (Figure 3). We utilized the same modest tolerance parameters in Truvari for benchmarking.

**Figure 3.**
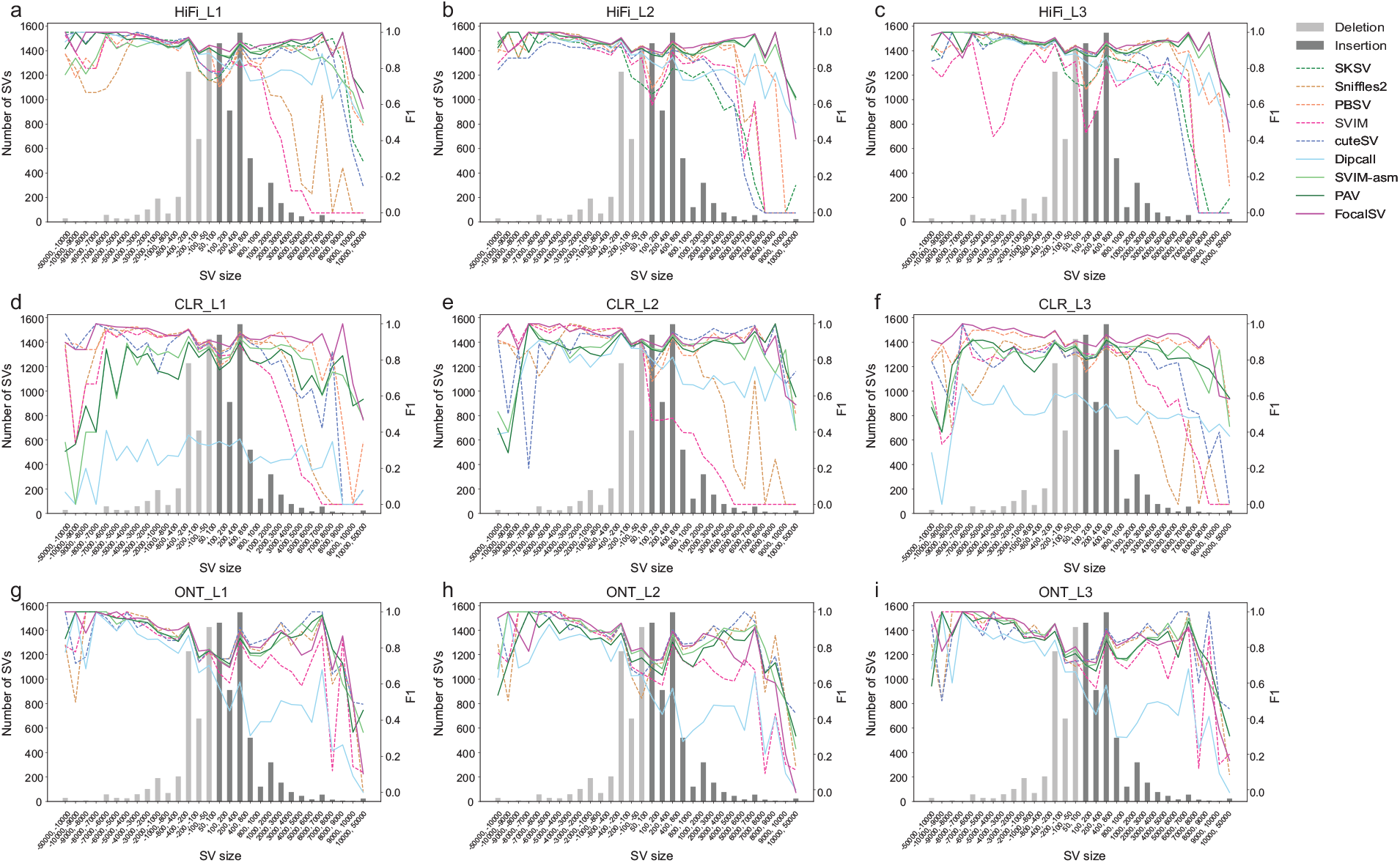
F1 accuracy of SV detection across various size ranges on nine long-read datasets. **(a-c)** F1 accuracy plot for three Hifi datasets. Negative ranges denote deletions and the positive ranges denote insertions. The bar plot illustrates the benchmark SV distribution across these size ranges. The line plot displays the F1 scores for four distinct detection methods. Dashed lines indicate alignment-based, while solid lines reprensent assembly-based methods. **(d-f)** F1 accuracy plot for three CLR datasets. **(g-i)** F1 accuracy plot for three ONT datasets.

Overall, assembly-based tools such as FocalSV, PAV, SVIM-asm, and Dipcall showed better resilience than alignment tools such as SKSV, Sniffles2, PBSV, SVIM, and cuteSV when processing HiFi data. Most alignment-based tools had a significant drop in performance with large INSs in the range of 2kb-50kb, while most assembly-based tools typically encountered performance challenges only with large INSs in the range of 9kb-50kb. Among all assembly-based tools, FocalSV continued to distinguish itself with exceptional performance across nearly all size ranges, except for 8kb-10kb deletions and 10kb-50kb insertions.

On CLR data, assembly-based tools except Dipcall still outperformed alignment-based tools in terms of resilience, especially for INSs. Across all size ranges, FocalSV maintained a top-tier performance. Notably, FocalSV’s performance is remarkably better than other tools on CLR L3, where it had the highest F1 score across all size ranges except for INSs in the range of 9kb-10kb and showed the least fluctuation.

On ONT data, the difference between alignment-based and assembly-based tools was less pronounced. Two alignment-based tools, cuteSV and Sniffles2, demonstrated resilience comparable to assembly-based tools. FocalSV emerged as a top-performing tool for small to large-sized deletions (50bp-8kb) and insertions (50bp-2kb). For large deletions (8kb-50kb) and insertions (2kb-50kb), FocalSV remained within the range of the best tools.

### FocalSV achieves robust SV detection across evaluation parameters

While assessing different SV callers against a gold standard, we recognized the potential impact of breakpoint shifts and sequence similarity issues on the evaluation. This acknowledgment arises from the fact that SVs often span substantial genomic regions. In previous evaluations, we selected a set of moderate-tolerance but fixed parameters to demonstrate the overall performance for each tool. However, the choice of parameters such as breakpoint shift tolerance and sequence similarity for identifying a call as a true positive, varies subjectively. Therefore, to comprehensively evaluate the effectiveness and stability of SV callers, we adjusted key evaluation parameters (p, P, r, and O) using Truvari for benchmarking [27]. Specifically, we systematically varied the range of four parameters, performing grid search benchmarking experiments using all parameter combinations to calculate the F1 scores for each tool. Parameters p, P, and O were adjusted in increments of 0.1 from 0 to 1, while r was adjusted in 100bp increments from 0 to 1000bp. Our goal was to examine how different levels of stringency or leniency in these parameters might impact SV detection across the nine datasets. We expected FocalSV to demonstrate relatively consistent and stable performance, even under more stringent conditions.

We initially evaluated deletion calls on Hifi L1 (Figure 4a). Based on the comprehensive benchmarking study by Liu et al. [8], which thoroughly examined the impact of various parameter combinations, such as p-O, P-r, O-r, p-P, p-r, and P-O, on deletion evaluation, we selected p and O as the representative parameter pair, as they had the most substantial influence on deletion performance. As shown in Figure 4, increasing the values of p and O imposed stricter correspondence requirements between the SV call and the gold standard, leading to a decline in F1 scores. Liu et al.’s heatmap and gradient analysis demonstrated distinct performance patterns among tools as more stringent thresholds were applied. They showed that all read alignment-based SV callers, except for pbsv, exhibited significant performance drops when p or O exceeded 0.7, with F1 scores falling below 5% when either parameter reached 1.0. In contrast, FocalSV and other assembly-based SV callers from Liu et al.’s study, namely Dipcall, SVIM-asm, and PAV, along with pbsv, maintained stable performance across the parameter grid. Even under the strictest conditions (p = 1.0, O = 1.0), these tools achieved F1 scores greater than 69%. Notably, under this strict exact-match criterion in terms of sequence similarity and reciprocal overlap ratio, FocalSV excelled with an exceptional F1 score above 74% (Figure 4a), outperforming other stable tools and demonstrating its remarkable robustness. Across all parameter settings, FocalSV consistently achieved higher F1 scores than Dipcall, SVIM-asm, PAV, and pbsv, demonstrating superior performance across the grid.

**Figure 4.**
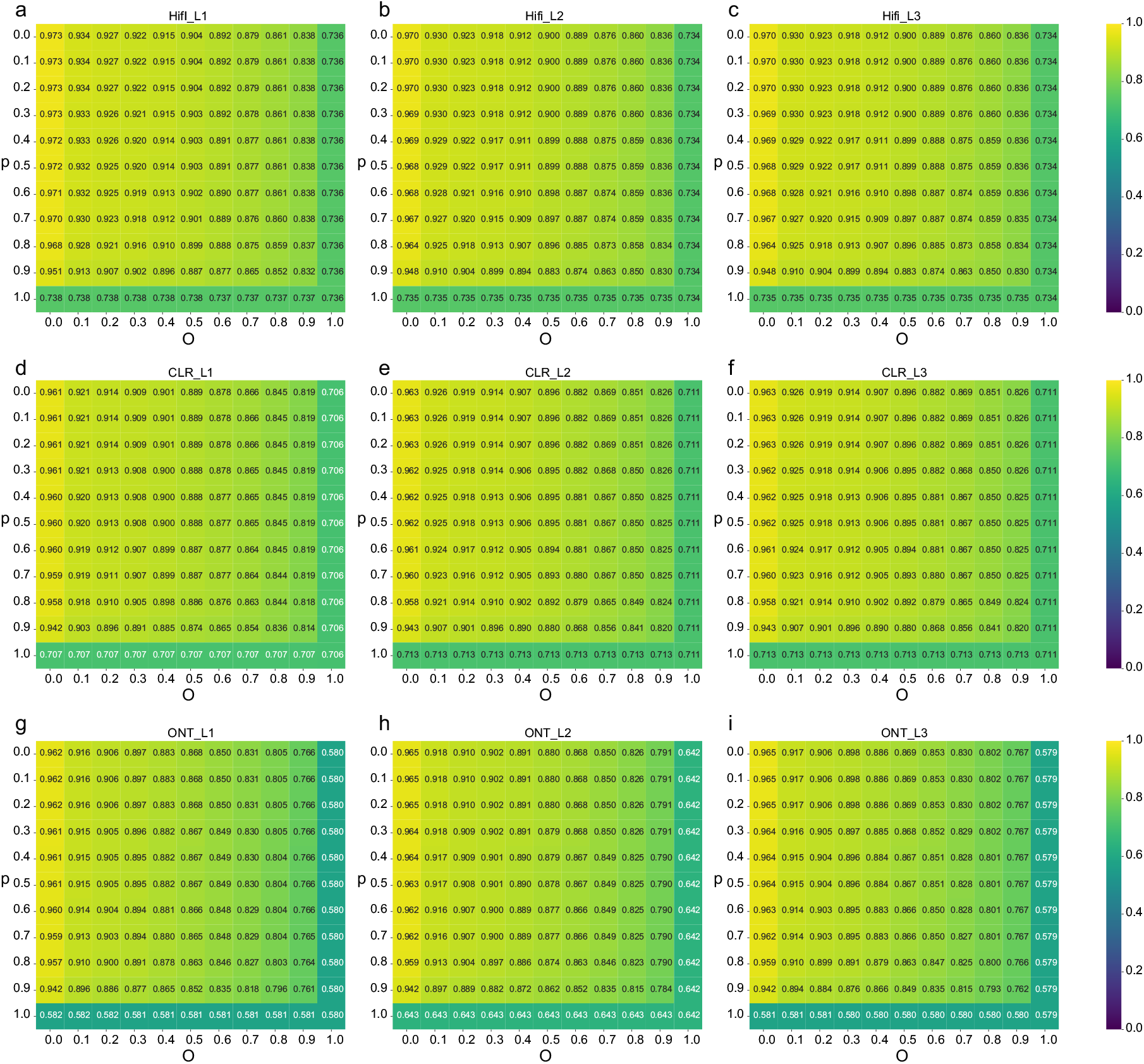
Deletion F1 accuracy by tuning different evaluation parameters (p and O). *O* is the minimum reciprocal overlap between SV call and gold standard SV. *p* is the minimum percentage of allele sequence similarity between SV call and gold standard SV. *O* and *p* vary from 0–1 with a 0.1 interval. **(a-c)** The F1 heatmap for deletions by FocalSV on three Hifi datasets. Every cell in the heatmap represents the F1 score under a specific pair of p and O evaluation. **(d-f)** The F1 heatmap for deletions by FocalSV on three CLR datasets. **(g-i)** The F1 heatmap for deletions by FocalSV on three ONT datasets.

For insertion evaluations on Hifi L1 (Figure 5a), the pair p and r were similarly selected as the most representative parameters based on Liu et al.’s findings [8]. Their study revealed that insertion detection was generally more sensitive to parameter variations compared to deletions. Combining their results and our experiment, we found that most tools, including FocalSV, experienced a decline in F1 scores when p exceeded

**Figure 5.**
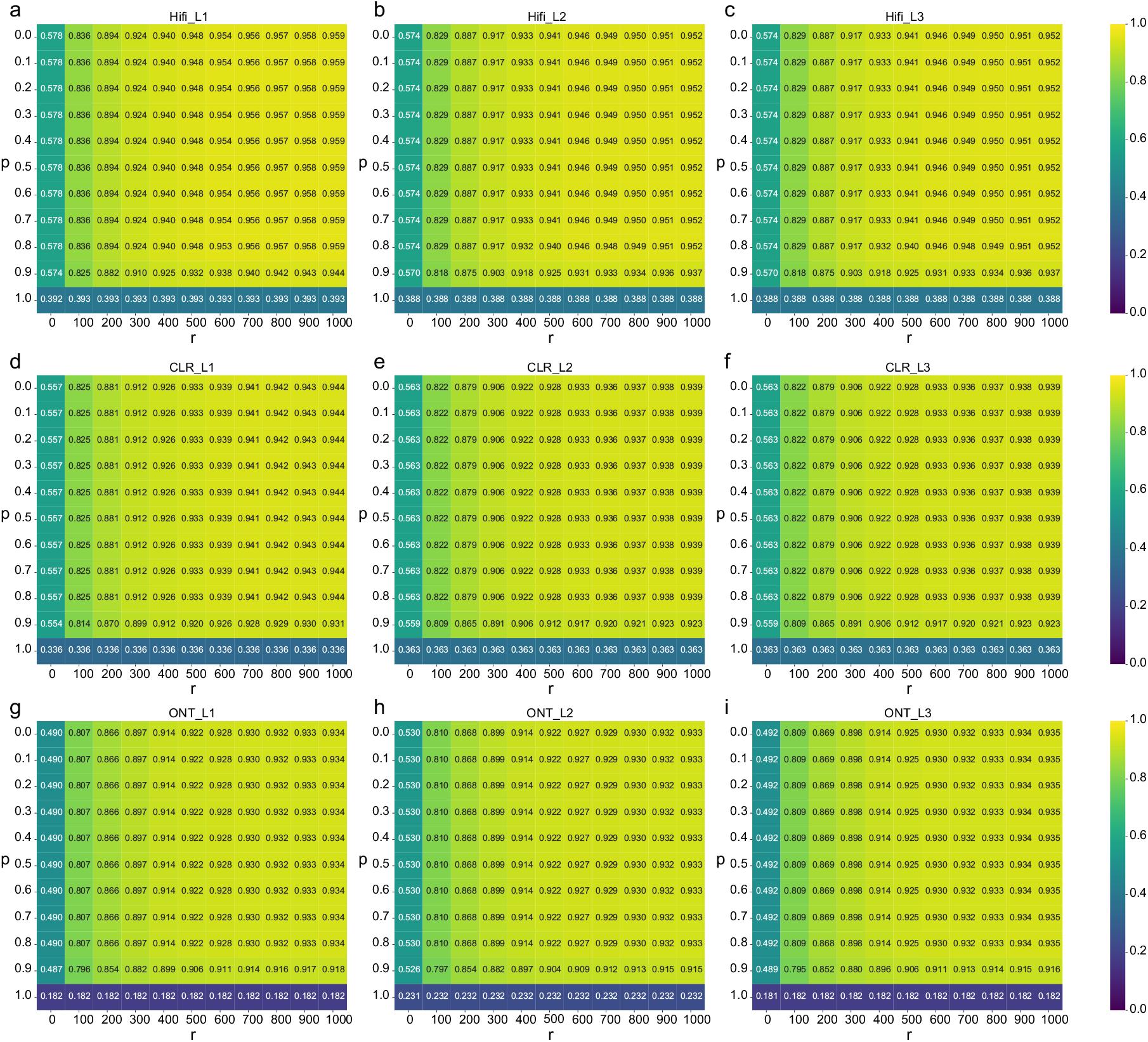
Insertion F1 accuracy by tuning different evaluation parameters (p and r). *r* is the maximum reference location distance between SV call and gold standard SV. *p* is the minimum percentage of allele sequence similarity between SV call and gold standard SV. *p* vary from 0–1 with a 0.1 interval. *r* varies from 0–1000 bp with a 100 bp interval. **(a-c)** The F1 heatmap for insertions by FocalSV on three Hifi datasets. Every cell in the heatmap represents the F1 score under a specific pair of p and r evaluation. **(d-f)** The F1 heatmap for insertions by FocalSV on three CLR datasets. **(g-i)** The F1 heatmap for insertions by FocalSV on three ONT datasets.

0.5 (i.e. when the required allele sequence similarity between the called SV and the benchmark exceeded 50%) (Figure 5a). Additionally, F1 scores decreased when r was reduced. Once r dropped to 200bp or less, indicating a more stringent reference distance threshold, performance deteriorated significantly across all tools. Similar to the trends observed for deletions, FocalSV, other assembly-based tools, and pbsv displayed greater robustness to stringent parameters compared to read alignment-based tools. Among all robust performing tools, FocalSV consistently achieved higher F1 scores than Dipcall, SVIM-asm, PAV, and pbsv, showing outstanding performance over the grid. We have also observed similar patterns on grid searches for FocalSV on other libraries (Figure 4b-i and Figure 5b-i). The robustness of assembly-based tools to evaluation parameters reflects their ability to accurately detect SV breakpoints and alternative allele sequences, as highlighted by Liu et al. In line with these findings, our results confirm that FocalSV, like other assembly-based tools, demonstrates strong resilience across various parameter settings, underscoring its capacity to accurately capture both SV breakpoints and alternative alleles. Notably, FocalSV consistently outperformed other assembly-based tools, including Dipcall, SVIM-asm, PAV, across all parameter configurations, further emphasizing its superior accuracy and reliability.

### FocalSV achieves superior performance in complex somatic SV detection in cancer data

To expand the evaluation to include complex SV detection, such as translocations (TRA), inversions (INV), and duplications (DUP), which involve various combinations of DNA rearrangements beyond deletions and insertions, we applied FocalSV and other relevant tools to detect complex somatic SVs in two publicly available cancer libraries [28]. Dipcall was excluded from this analysis, as it was not designed to detect complex SVs. Although PAV is designed to detect complex SVs like inversions, it failed to detect any inversions in the cancer data and was therefore also excluded from this analysis.

We conducted a comparative analysis by applying the selected tools to two publicly available tumor-normal paired libraries (Pacbio CLR and ONT), as provided by Talsania et al [16]. These libraries, along with the high-confidence HCC1395 somatic SV call set serving as the benchmark gold standard, formed the basis for evaluating the detection of three classes of somatic complex SVs. Each assembly-based tool was first applied independently to each library, generating VCF files. These VCFs were then processed using SURVIVOR [17] to identify somatic variants by comparing paired normal-tumor VCFs. The identified somatic SVs were subsequently compared to the gold standard, which contained 1777 SVs, including 551 insertions, 717 deletions, 146 translocations, 133 inversions, and 230 duplications. Given the incompleteness of the high-confidence call set, our evaluation primarily focused on recall.

FocalSV consistently outperformed other tools in recall across all libraries and complex SV types, particularly excelling in the detection of translocations and duplications. Overall, FocalSV achieved the highest recall for all complex SV types on ONT data. On Pacbio CLR data, FocalSV led in recall for translocations and duplications, and achieved second-highest recall for inversions. Specifically, on ONT data, FocalSV outperformed the second-ranked tool by 26%, 11%, and 24% for translocations, inversions, and duplications, respectively. Similarly, on CLR data, FocalSV’s recall exceeded the second-ranked tool by 13% for translocations and 22% for duplications. For inversions, FocalSV ranked second by a small margin of 1% compared to pbsv, but still performed substantially better than the third-ranked tool, with an 8% difference.

### FocalSV achieves highly efficient region-based SV detection by optimized computational resource allocation

Finally, we evaluated FocalSV’s memory requirement and runtime performance. For a relatively large SV - an insertion of 12.6kb on chromosome 21 - FocalSV finished execution in 32 seconds of elapsed time and 5 minutes and 20 seconds of CPU time, using a maximum 5.6MB of memory when run with 8 threads. In comparison, detecting the smallest SV - a deletion of 50bp on chromosome 21 - took a similar elapsed time of 33 seconds and required 5 minutes and 30 seconds for CPU time, with a higher memory usage of 0.76GB. On average, the CPU time across these tasks was approximately 5.5 minutes, reflecting consistent computational effort regardless of SV size.

FocalSV is designed to support multi-level parallelization, allowing tasks to be efficiently distributed across multiple servers, with each server assigned to distinct genomic regions. Within each region, FocalSV utilizes multithreading to manage assembly and variant detection tasks, maximizing CPU core utilization. This design ensures high efficiency in large-scale SV analysis.

## Discussion

Through comprehensive benchmarking analysis, FocalSV exhibited strong performance across various SV types when applied to PacBio data, surpassing other tools in overall accuracy. On ONT data, FocalSV demonstrated state-of-the-art performance for deletion (DEL) calls and maintained competitive accuracy for insertion (INS) calls. Furthermore, FocalSV achieved the highest genotype accuracy among assembly-based tools across different data types and SV categories, highlighting its robustness and reliability.

FocalSV’s strong performance is driven by its comprehensive algorithm. FocalSV integrates Longshot for phasing, using its results to guide phased local assembly and generate accurate haplotype-resolved contigs. It further enhances traditional read-based signature extraction methodologies by incorporating a contig-based signature extraction technique to construct the draft SV map. To improve accuracy, FocalSV implements a strategy that revisits the read-based BAM file to extract additional signatures, enabling the filtering of false positives and the correction of false genotypes. These methodological advancements contribute to FocalSV’s effectiveness as an SV detection tool.

FocalSV introduces an innovative and efficient strategy for detecting complex SVs. For large insertions and deletions, FocalSV builds upon VolcanoSV’s established methodology, adapting it for region-specific analysis. While VolcanoSV relies on a comprehensive whole-genome assembly approach for large INDEL detection, FocalSV focuses on short, localized assemblies within targeted regions. This regional approach, while efficient, may miss certain SV-related signals due to its limited assembly scope. As a result, we did not directly compare the large INDEL detection performance of FocalSV and VolcanoSV in all table results, acknowledging the inherent differences in their strategies. Overall, FocalSV demonstrated comparable performance to VolcanoSV across all libraries, maintaining similar average F1 scores and recall. Specifically, for insertions, FocalSV achieved a slightly higher average F1 score, outperforming VolcanoSV by 0.56% due to improved precision. For deletions, FocalSV’s average F1 score was 0.36% lower than VolcanoSV’s, attributed to a slight decrease in precision. Both tools exhibited nearly identical genotype concordance, with FocalSV trailing VolcanoSV by just 0.14% for insertions and 0.08% for deletions - differences that are negligible. However, by avoiding the computational demands of whole-genome assembly, FocalSV’s region-based approach provides greater flexibility and efficiency, particularly for analyses concentrated on specific genomic regions, such as medically relevant loci or SV hotspots. For such targeted applications, FocalSV offers a streamlined and effective solution.

## Conclusions

FocalSV represents a notable advancement in SV detection for target regions or regions of interest, demonstrating strong performance on both PacBio and ONT data. Its region-specific design, combined with advanced algorithmic strategies, provides a highly efficient and accurate solution for detecting diverse SV types across different sequencing platforms.

## Data availability

PacBio CLR, Hifi, and ONT sequencing reads for HG002 are available at GIAB and NCBI. The high-confidence HCC1395 somatic SV callset and the Pacbio and ONT Tumor-Normal paired libraries of HCC1395 are publicly accessible at NCBI. Table 1 lists hyperlinks for all 13 previously mentioned real datasets. The Tier1 benchmark SV callset and high-confidence HG002 region were obtained from https://ftp-trace.ncbi.nlm.nih.gov/ReferenceSamples/giab/data/AshkenazimTrio/analysis/NIST_SVs_Integration_v0.6/. All VCF files that support the findings of this study are available from https://doi.org/10.5281/zenodo.14187482.

## Code availability

FocalSV can be found at https://github.com/maiziezhoulab/FocalSV.

## Competing interests

The authors declare that they have no competing interests.

## Author’s contributions

X.M.Z. conceived and led this work. C.L., Z.Z., and X.M.Z. designed the framework. C.L. and Z.Z. implemented the framework and performed analyses. C.L., Z.Z., and X.M.Z. wrote the manuscript.

## Acknowledgements

This work was supported by the NIH NIGMS Maximizing Investigators’ Research Award (MIRA) R35 GM146960.

## Notes

### Competing Interest Statement

The authors have declared no competing interest.

