## Supplementary file for "FocalSV: target region-based structural variant assembly and refinement using single-molecule long read sequencing data"

**Supplementary References** **10**

### 1 Supplementary Tables

Supplementary Table S1: Deletion and Insertion SV calling results for HIFI Lib1-3 from 9 tools

| **DEL** | | **FocalSV** | **PAV** | **SVIM-asm** | **Dipcall** | **cuteSV** | **SVIM** | **PBSV** | **Sniffles2** | **SKSV** |
|---|---|---|---|---|---|---|---|---|---|---|
| | True Positive (TP) | 3925 | 3837 | 3865 | 3812 | 3641 | 3654 | 3846 | 3484 | 3509 |
| | False Positive (FP) | 289 | 321 | 350 | 408 | 487 | 530 | 288 | 676 | 605 |
| | False Negative (FN) | 191 | 279 | 251 | 304 | 475 | 462 | 270 | 632 | 607 |
| | Precision | **0.931** | 0.923 | 0.917 | 0.903 | 0.882 | 0.873 | 0.930 | 0.838 | 0.853 |
| **Hifi_L1** | Recall | **0.954** | 0.932 | 0.939 | 0.926 | 0.885 | 0.888 | 0.934 | 0.847 | 0.853 |
| | F1 | **0.942** | 0.928 | 0.928 | 0.915 | 0.883 | 0.881 | 0.932 | 0.842 | 0.853 |
| | Genotype TP | 3782 | 3726 | 3782 | 3688 | 3607 | 3607 | 3758 | 3364 | 3438 |
| | Genotype FP | 143 | 111 | 83 | 124 | 34 | 47 | 88 | 120 | 71 |
| | Genotype Concordance | **0.991** | 0.971 | 0.979 | 0.968 | **0.991** | 0.987 | 0.977 | 0.966 | 0.980 |
| | True Positive (TP) | 3910 | 3874 | 3892 | 3818 | 3586 | 3926 | 3860 | 3904 | 3248 |
| | False Positive (FP) | 310 | 335 | 338 | 419 | 236 | 809 | 326 | 309 | 537 |
| | False Negative (FN) | 206 | 242 | 224 | 298 | 530 | 190 | 256 | 212 | 868 |
| | Precision | 0.927 | 0.920 | 0.920 | 0.901 | **0.938** | 0.829 | 0.922 | 0.927 | 0.858 |
| **Hifi_L2** | Recall | 0.950 | 0.941 | 0.946 | 0.928 | 0.871 | **0.954** | 0.938 | 0.949 | 0.789 |
| | F1 | **0.938** | 0.931 | 0.933 | 0.914 | 0.904 | 0.887 | 0.930 | 0.937 | 0.822 |
| | Genotype TP | 3871 | 3768 | 3838 | 3698 | 3495 | 3844 | 3790 | 3838 | 3166 |
| | Genotype FP | 39 | 106 | 54 | 120 | 91 | 82 | 70 | 66 | 82 |
| | Genotype Concordance | **0.990** | 0.973 | 0.986 | 0.969 | 0.975 | 0.979 | 0.982 | 0.983 | 0.975 |
| | True Positive (TP) | 3919 | 3848 | 3887 | 3818 | 3722 | 3924 | 3866 | 3911 | 3352 |
| | False Positive (FP) | 310 | 348 | 352 | 427 | 228 | 1287 | 314 | 303 | 551 |
| | False Negative (FN) | 197 | 268 | 229 | 298 | 394 | 192 | 250 | 205 | 764 |
| | Precision | 0.927 | 0.917 | 0.917 | 0.899 | **0.942** | 0.753 | 0.925 | 0.928 | 0.859 |
| **Hifi_L3** | Recall | 0.952 | 0.935 | 0.944 | 0.928 | 0.904 | **0.953** | 0.939 | 0.950 | 0.814 |
| | F1 | **0.939** | 0.926 | 0.931 | 0.913 | 0.923 | 0.841 | 0.932 | 0.939 | 0.836 |
| | Genotype TP | 3879 | 3718 | 3832 | 3697 | 3649 | 3859 | 3812 | 3851 | 3287 |
| | Genotype FP | 40 | 130 | 55 | 121 | 73 | 65 | 54 | 60 | 65 |
| | Genotype Concordance | **0.990** | 0.966 | 0.986 | 0.968 | 0.980 | 0.983 | 0.986 | 0.985 | 0.981 |

| **INS** | | FocalSV | PAV | SVIM-asm | Dipcall | cuteSV | SVIM | PBSV | **Sniffles2** | **SKSV** |
|---|---|---|---|---|---|---|---|---|---|---|
| | True Positive (TP) | 4961 | 4917 | 4916 | 4824 | 4348 | 3677 | 4088 | 4053 | 4572 |
| | False Positive (FP) | 501 | 778 | 766 | 1801 | 620 | 645 | 455 | 1075 | 736 |
| | False Negative (FN) | 320 | 364 | 365 | 457 | 933 | 1604 | 1193 | 1228 | 709 |
| | Precision | **0.908** | 0.863 | 0.865 | 0.728 | 0.875 | 0.851 | 0.900 | 0.790 | 0.861 |
| **Hifi_L1** | Recall | **0.939** | 0.931 | 0.931 | 0.914 | 0.823 | 0.696 | 0.774 | 0.768 | 0.866 |
| | F1 | **0.924** | 0.896 | 0.897 | 0.810 | 0.849 | 0.766 | 0.832 | 0.779 | 0.864 |
| | Genotype TP | 4866 | 4699 | 4809 | 3755 | 4307 | 3309 | 3586 | 3621 | 4495 |
| | Genotype FP | 95 | 218 | 107 | 1069 | 41 | 368 | 502 | 432 | 77 |
| | Genotype Concordance | 0.981 | 0.956 | 0.978 | 0.778 | **0.991** | 0.900 | 0.877 | 0.893 | 0.983 |
| | True Positive (TP) | 4952 | 4929 | 4907 | 4837 | 4351 | 4838 | 4050 | 4849 | 3257 |
| | False Positive (FP) | 563 | 779 | 774 | 1776 | 391 | 2976 | 457 | 642 | 522 |
| | False Negative (FN) | 329 | 352 | 374 | 444 | 930 | 443 | 1231 | 432 | 2024 |
| | Precision | 0.898 | 0.864 | 0.864 | 0.731 | **0.918** | 0.619 | 0.899 | 0.883 | 0.862 |
| **Hifi_L2** | Recall | **0.938** | 0.933 | 0.929 | 0.916 | 0.824 | 0.916 | 0.767 | 0.918 | 0.617 |
| | F1 | **0.917** | 0.897 | 0.895 | 0.813 | 0.868 | 0.739 | 0.828 | 0.900 | 0.719 |
| | Genotype TP | 4868 | 4737 | 4793 | 3798 | 4248 | 4546 | 3864 | 4703 | 3182 |
| | Genotype FP | 84 | 192 | 114 | 1039 | 103 | 292 | 186 | 146 | 75 |
| | Genotype Concordance | **0.983** | 0.961 | 0.977 | 0.785 | 0.976 | 0.940 | 0.954 | 0.970 | 0.977 |
| | True Positive (TP) | 4953 | 4912 | 4919 | 4846 | 4539 | 4866 | 4048 | 4901 | 3751 |
| | False Positive (FP) | 574 | 781 | 777 | 1795 | 415 | 5940 | 499 | 628 | 556 |
| | False Negative (FN) | 328 | 369 | 362 | 435 | 742 | 415 | 1233 | 380 | 1530 |
| | Precision | 0.896 | 0.863 | 0.864 | 0.730 | **0.916** | 0.450 | 0.890 | 0.886 | 0.871 |
| **Hifi_L3** | Recall | **0.938** | 0.930 | 0.932 | 0.918 | 0.860 | 0.921 | 0.767 | 0.928 | 0.710 |
| | F1 | **0.917** | 0.895 | 0.896 | 0.813 | 0.887 | 0.605 | 0.824 | 0.907 | 0.782 |
| | Genotype TP | 4878 | 4661 | 4818 | 3794 | 4466 | 4586 | 3894 | 4757 | 3699 |
| | Genotype FP | 75 | 251 | 101 | 1052 | 73 | 280 | 154 | 144 | 52 |
| | Genotype Concordance | 0.985 | 0.949 | 0.980 | 0.783 | 0.984 | 0.943 | 0.962 | 0.971 | **0.986** |

Supplementary Table S2: Deletion and Insertion SV calling results for CLR Lib1-3 from 8 tools

| **DEL** | | FocalSV | PAV | SVIM-asm | Dipcall | cuteSV | SVIM | PBSV | Sniffles2 |
|---|---|---|---|---|---|---|---|---|---|
| **CLR_L1** | True Positive (TP) | 3876 | 3576 | 3544 | 866 | 3679 | 3774 | 3782 | 3779 |
| | False Positive (FP) | 348 | 811 | 460 | 185 | 318 | 339 | 242 | 307 |
| | False Negative (FN) | 240 | 540 | 572 | 3250 | 437 | 342 | 334 | 337 |
| | Precision | 0.918 | 0.815 | 0.885 | 0.824 | 0.920 | 0.918 | **0.940** | 0.925 |
| | Recall | **0.942** | 0.869 | 0.861 | 0.210 | 0.894 | 0.917 | 0.919 | 0.918 |
| | F1 | **0.930** | 0.841 | 0.873 | 0.335 | 0.907 | 0.917 | 0.929 | 0.922 |
| | Genotype TP | 3828 | 3433 | 3375 | 740 | 3629 | 3694 | 3655 | 3729 |
| | Genotype FP | 48 | 143 | 169 | 126 | 50 | 80 | 127 | 50 |
| | Genotype Concordance | **0.988** | 0.960 | 0.952 | 0.855 | 0.986 | 0.979 | 0.966 | 0.987 |
| **CLR_L2** | True Positive (TP) | 3903 | 3664 | 3775 | 3579 | 3786 | 3863 | 3826 | 3796 |
| | False Positive (FP) | 338 | 353 | 328 | 451 | 355 | 273 | 275 | 289 |
| | False Negative (FN) | 213 | 452 | 341 | 537 | 330 | 253 | 290 | 320 |
| | Precision | 0.920 | 0.912 | 0.920 | 0.888 | 0.914 | **0.934** | 0.933 | 0.929 |
| | Recall | **0.948** | 0.890 | 0.917 | 0.870 | 0.920 | 0.939 | 0.930 | 0.922 |
| | F1 | 0.934 | 0.901 | 0.919 | 0.879 | 0.917 | **0.936** | 0.931 | 0.926 |
| | Genotype TP | 3853 | 3605 | 3717 | 3377 | 3759 | 3808 | 3742 | 3696 |
| | Genotype FP | 50 | 59 | 58 | 202 | 27 | 55 | 84 | 100 |
| | Genotype Concordance | 0.987 | 0.984 | 0.985 | 0.944 | **0.993** | 0.986 | 0.978 | 0.974 |
| **CLR_L3** | True Positive (TP) | 3809 | 3472 | 3469 | 1907 | 3297 | 3265 | 3758 | 3224 |
| | False Positive (FP) | 311 | 459 | 360 | 389 | 278 | 245 | 302 | 203 |
| | False Negative (FN) | 307 | 644 | 647 | 2209 | 819 | 851 | 358 | 892 |
| | Precision | 0.925 | 0.883 | 0.906 | 0.831 | 0.922 | 0.930 | 0.926 | **0.941** |
| | Recall | **0.925** | 0.844 | 0.843 | 0.463 | 0.801 | 0.793 | 0.913 | 0.783 |
| | F1 | **0.925** | 0.863 | 0.873 | 0.595 | 0.857 | 0.856 | 0.919 | 0.855 |
| | Genotype TP | 3704 | 3321 | 3336 | 1607 | 3223 | 3226 | 3637 | 3095 |
| | Genotype FP | 105 | 151 | 133 | 300 | 74 | 39 | 121 | 129 |
| | Genotype Concordance | 0.972 | 0.957 | 0.962 | 0.843 | **0.978** | 0.988 | 0.968 | 0.960 |

| **INS** | | FocalSV | PAV | SVIM-asm | Dipcall | cuteSV | SVIM | PBSV | Sniffles2 |
|---|---|---|---|---|---|---|---|---|---|
| **CLR_L1** | True Positive (TP) | 4903 | 4815 | 4750 | 1097 | 4548 | 3760 | 4213 | 4576 |
| | False Positive (FP) | 597 | 1868 | 1172 | 479 | 693 | 295 | 310 | 623 |
| | False Negative (FN) | 378 | 466 | 531 | 4184 | 733 | 1521 | 1068 | 705 |
| | Precision | 0.892 | 0.721 | 0.802 | 0.696 | 0.868 | 0.927 | **0.932** | 0.880 |
| | Recall | **0.928** | 0.912 | 0.900 | 0.208 | 0.861 | 0.712 | 0.798 | 0.867 |
| | F1 | **0.910** | 0.805 | 0.848 | 0.320 | 0.865 | 0.806 | 0.859 | 0.873 |
| | Genotype TP | 4753 | 4418 | 4370 | 746 | 4459 | 2320 | 3689 | 4426 |
| | Genotype FP | 150 | 397 | 380 | 351 | 89 | 1440 | 524 | 150 |
| | Genotype Concordance | 0.969 | 0.918 | 0.920 | 0.680 | **0.980** | 0.617 | 0.876 | 0.967 |
| **CLR_L2** | True Positive (TP) | 4939 | 4846 | 4928 | 4654 | 4852 | 1460 | 4114 | 4272 |
| | False Positive (FP) | 675 | 913 | 896 | 2317 | 983 | 863 | 664 | 993 |
| | False Negative (FN) | 342 | 435 | 353 | 627 | 429 | 3821 | 1167 | 1009 |
| | Precision | **0.880** | 0.842 | 0.846 | 0.668 | 0.832 | 0.629 | 0.861 | 0.811 |
| | Recall | **0.935** | 0.918 | 0.933 | 0.881 | 0.919 | 0.277 | 0.779 | 0.809 |
| | F1 | **0.907** | 0.878 | 0.888 | 0.760 | 0.873 | 0.384 | 0.818 | 0.810 |
| | Genotype TP | 4792 | 4646 | 4769 | 3146 | 4802 | 644 | 3683 | 4057 |
| | Genotype FP | 147 | 200 | 159 | 1508 | 50 | 816 | 431 | 215 |
| | Genotype Concordance | 0.970 | 0.959 | 0.968 | 0.676 | **0.990** | 0.441 | 0.895 | 0.950 |
| **CLR_L3** | True Positive (TP) | 4829 | 4843 | 4774 | 2497 | 4216 | 3986 | 4221 | 3719 |
| | False Positive (FP) | 561 | 1294 | 1083 | 1637 | 566 | 362 | 599 | 436 |
| | False Negative (FN) | 452 | 438 | 507 | 2784 | 1065 | 1295 | 1060 | 1562 |
| | Precision | 0.896 | 0.789 | 0.815 | 0.604 | 0.882 | **0.917** | 0.876 | 0.895 |
| | Recall | 0.914 | **0.917** | 0.904 | 0.473 | 0.798 | 0.755 | 0.799 | 0.704 |
| | F1 | **0.905** | 0.848 | 0.857 | 0.530 | 0.838 | 0.828 | 0.836 | 0.788 |
| | Genotype TP | 4591 | 4322 | 4290 | 1355 | 4057 | 3495 | 3742 | 3445 |
| | Genotype FP | 238 | 521 | 484 | 1142 | 159 | 491 | 479 | 274 |
| | Genotype Concordance | 0.951 | 0.892 | 0.899 | 0.543 | **0.962** | 0.877 | 0.887 | 0.926 |

Supplementary Table S3: Deletion and Insertion SV calling results for ONT Lib1-3 from 7 tools

| **DEL** | | FocalSV | PAV | SVIM-asm | Dipcall | cuteSV | SVIM | Sniffles2 |
|---|---|---|---|---|---|---|---|---|
| **ONT_L1** | True Positive (TP) | 3865 | 3819 | 3865 | 3734 | 3819 | 3828 | 3805 |
| | False Positive (FP) | 341 | 373 | 389 | 584 | 363 | 384 | 349 |
| | False Negative (FN) | 251 | 297 | 251 | 382 | 297 | 288 | 311 |
| | Precision | **0.919** | 0.911 | 0.909 | 0.865 | 0.913 | 0.909 | 0.916 |
| | Recall | **0.939** | 0.928 | **0.939** | 0.907 | 0.928 | 0.930 | 0.924 |
| | F1 | **0.929** | 0.919 | 0.924 | 0.886 | 0.921 | 0.919 | 0.920 |
| | Genotype TP | 3824 | 3771 | 3818 | 3459 | 3784 | 3763 | 3774 |
| | Genotype FP | 41 | 48 | 47 | 275 | 35 | 65 | 31 |
| | Genotype Concordance | 0.989 | 0.987 | 0.988 | 0.926 | 0.991 | 0.983 | **0.992** |
| **ONT_L2** | True Positive (TP) | 3868 | 3665 | 3857 | 3647 | 3803 | 3809 | 3801 |
| | False Positive (FP) | 335 | 368 | 379 | 661 | 642 | 602 | 970 |
| | False Negative (FN) | 248 | 451 | 259 | 469 | 313 | 307 | 315 |
| | Precision | **0.920** | 0.909 | 0.911 | 0.847 | 0.856 | 0.864 | 0.797 |
| | Recall | **0.940** | 0.890 | 0.937 | 0.886 | 0.924 | 0.925 | 0.924 |
| | F1 | **0.930** | 0.900 | 0.924 | 0.866 | 0.888 | 0.893 | 0.855 |
| | Genotype TP | 3834 | 3618 | 3808 | 3289 | 3760 | 3734 | 3764 |
| | Genotype FP | 34 | 47 | 49 | 358 | 43 | 75 | 37 |
| | Genotype Concordance | **0.991** | 0.987 | 0.987 | 0.902 | 0.989 | 0.980 | 0.990 |
| **ONT_L3** | True Positive (TP) | 3853 | 3741 | 3857 | 3669 | 3804 | 3820 | 3804 |
| | False Positive (FP) | 323 | 372 | 380 | 628 | 478 | 480 | 516 |
| | False Negative (FN) | 263 | 375 | 259 | 447 | 312 | 296 | 312 |
| | Precision | **0.923** | 0.910 | 0.910 | 0.854 | 0.888 | 0.888 | 0.881 |
| | Recall | 0.936 | 0.909 | **0.937** | 0.891 | 0.924 | 0.928 | 0.924 |
| | F1 | **0.929** | 0.909 | 0.924 | 0.872 | 0.906 | 0.908 | 0.902 |
| | Genotype TP | 3819 | 3694 | 3811 | 3341 | 3770 | 3761 | 3778 |
| | Genotype FP | 34 | 47 | 46 | 328 | 34 | 59 | 26 |
| | Genotype Concordance | 0.991 | 0.987 | 0.988 | 0.911 | 0.991 | 0.985 | **0.993** |

| **INS** | | FocalSV | PAV | SVIM-asm | Dipcall | cuteSV | SVIM | Sniffles2 |
|---|---|---|---|---|---|---|---|---|
| **ONT_L1** | True Positive (TP) | 4903 | 4896 | 4929 | 4724 | 4835 | 4671 | 4813 |
| | False Positive (FP) | 773 | 795 | 851 | 2492 | 518 | 984 | 595 |
| | False Negative (FN) | 378 | 385 | 352 | 557 | 446 | 610 | 468 |
| | Precision | 0.864 | 0.860 | 0.853 | 0.655 | **0.903** | 0.826 | 0.890 |
| | Recall | 0.928 | 0.927 | **0.933** | 0.895 | 0.916 | 0.885 | 0.911 |
| | F1 | 0.895 | 0.893 | 0.891 | 0.756 | **0.909** | 0.854 | 0.901 |
| | Genotype TP | 4812 | 4760 | 4840 | 3011 | 4781 | 4003 | 4618 |
| | Genotype FP | 91 | 136 | 89 | 1713 | 54 | 668 | 195 |
| | Genotype Concordance | 0.981 | 0.972 | 0.982 | 0.637 | **0.989** | 0.857 | 0.960 |
| **ONT_L2** | True Positive (TP) | 4943 | 4713 | 4906 | 4584 | 4783 | 4587 | 4763 |
| | False Positive (FP) | 789 | 850 | 862 | 2598 | 540 | 1031 | 625 |
| | False Negative (FN) | 338 | 568 | 375 | 697 | 498 | 694 | 518 |
| | Precision | 0.862 | 0.847 | 0.851 | 0.638 | **0.899** | 0.8165 | 0.884 |
| | Recall | **0.936** | 0.892 | 0.929 | 0.868 | 0.906 | 0.8686 | 0.902 |
| | F1 | 0.898 | 0.869 | 0.888 | 0.736 | **0.902** | 0.8417 | 0.893 |
| | Genotype TP | 4859 | 4580 | 4804 | 2716 | 4711 | 3785 | 4538 |
| | Genotype FP | 84 | 133 | 102 | 1868 | 72 | 802 | 225 |
| | Genotype Concordance | 0.983 | 0.972 | 0.979 | 0.593 | **0.985** | 0.8252 | 0.953 |
| **ONT_L3** | True Positive (TP) | 4911 | 4791 | 4894 | 4594 | 4809 | 4693 | 4790 |
| | False Positive (FP) | 746 | 860 | 899 | 2537 | 520 | 1062 | 613 |
| | False Negative (FN) | 370 | 490 | 387 | 687 | 472 | 588 | 491 |
| | Precision | 0.868 | 0.848 | 0.845 | 0.644 | **0.902** | 0.816 | 0.887 |
| | Recall | **0.930** | 0.907 | 0.927 | 0.870 | 0.911 | 0.889 | 0.907 |
| | F1 | 0.898 | 0.877 | 0.884 | 0.740 | **0.907** | 0.851 | 0.897 |
| | Genotype TP | 4831 | 4652 | 4801 | 2783 | 4759 | 4081 | 4581 |
| | Genotype FP | 80 | 139 | 93 | 1811 | 50 | 612 | 209 |
| | Genotype Concordance | 0.984 | 0.971 | 0.981 | 0.606 | **0.990** | 0.870 | 0.956 |

### 2 Supplementary Figures

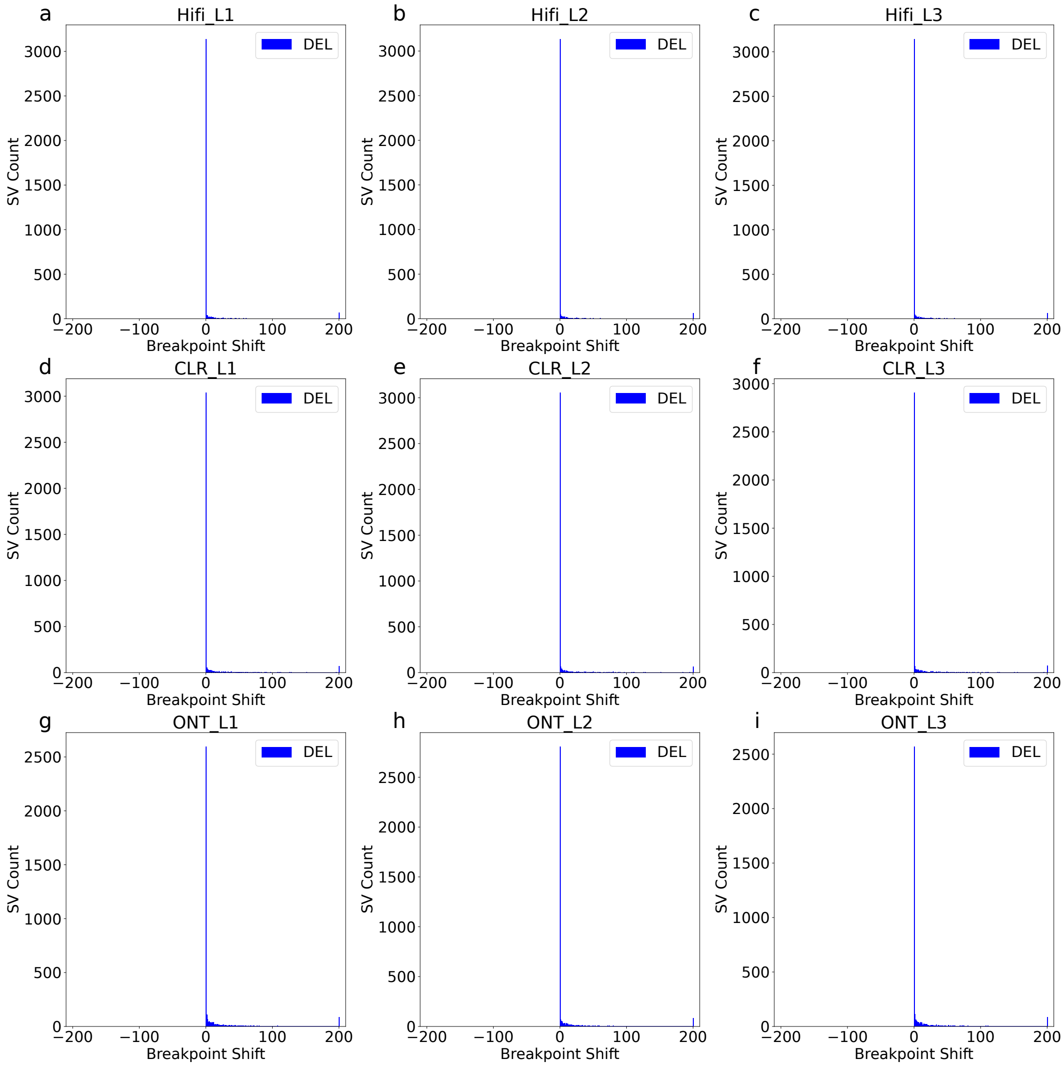

Supplementary Figure 1: **Distribution of breakpoint shift for deletion SVs across all libraries. (a-c)** Distribution of breakpoint shift for Hifi datasets. **(d-f)** Distribution of breakpoint shift for CLR datasets. **(g-i)** Distribution of breakpoint shift for ONT datasets. The evaluation was done by moderate Truvari parameters ($p$=0.5, $P$=0.5, $O$=0.01, $r$=500)

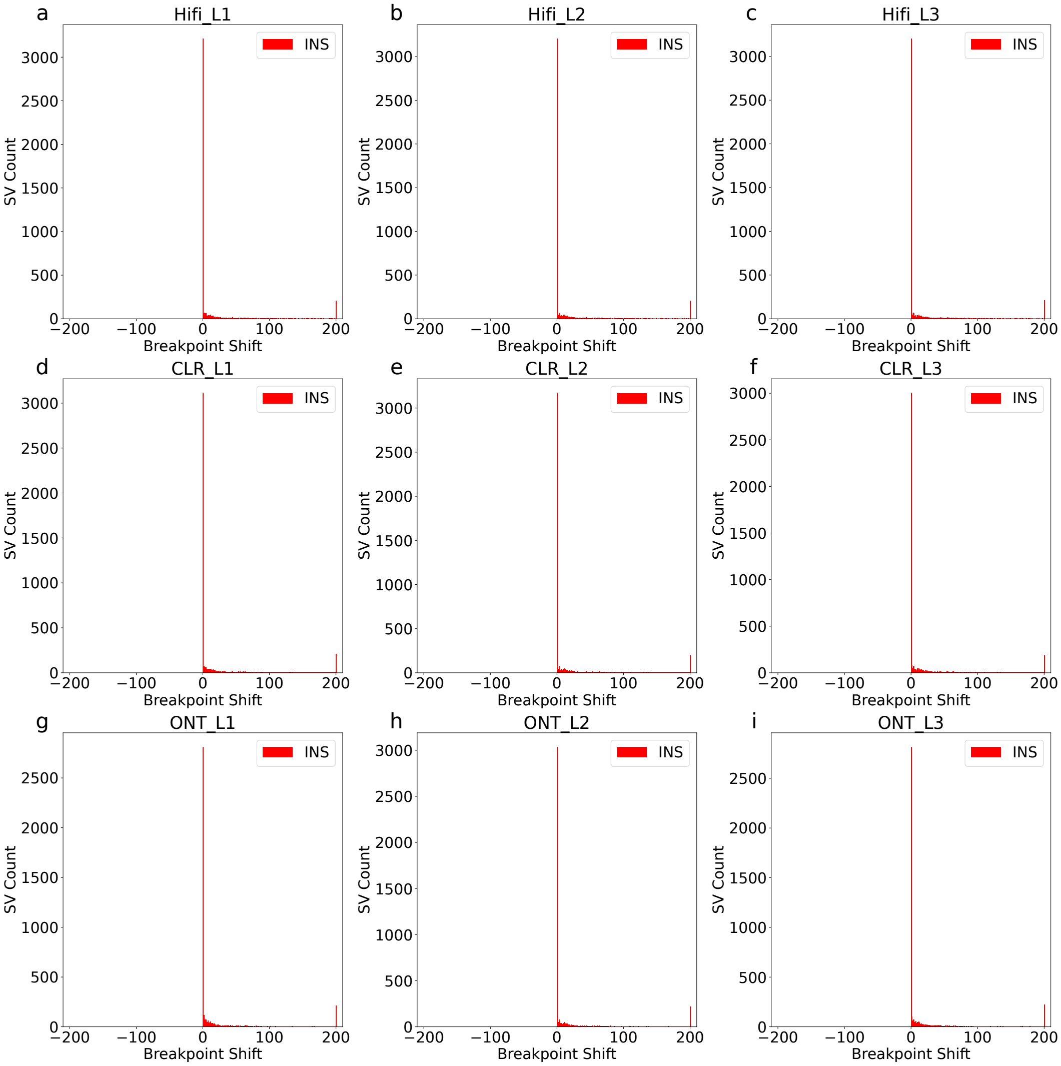

Supplementary Figure 2: **Distribution of breakpoint shift for insertion SVs across all libraries. (a-c)** Distribution of breakpoint shift for Hifi datasets. **(d-f)** Distribution of breakpoint shift for CLR datasets. **(g-i)** Distribution of breakpoint shift for ONT datasets. The evaluation was done by moderate Truvari parameters ($p$=0.5, $P$=0.5, $O$=0.01, $r$=500)

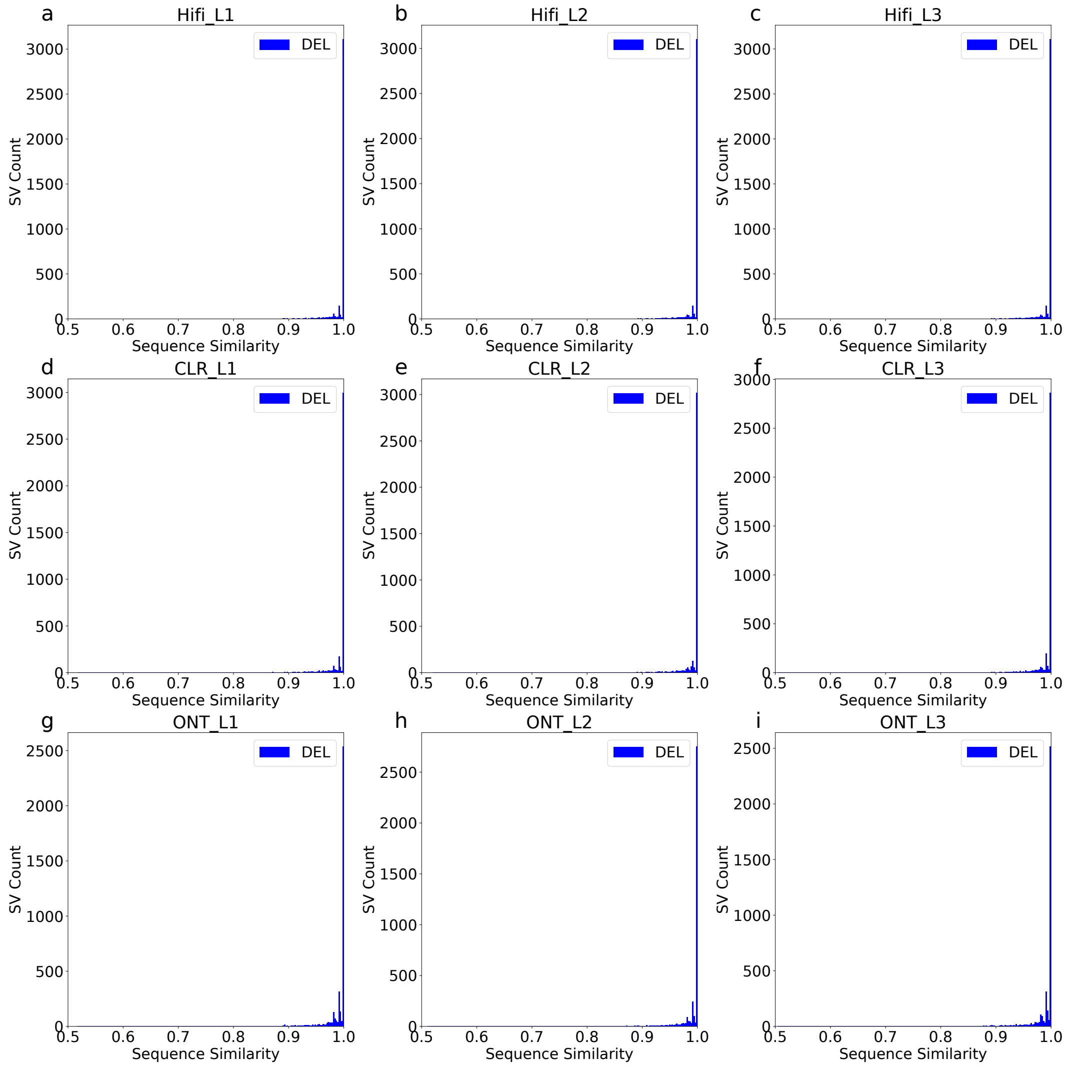

Supplementary Figure 3: **Distribution of sequence similarity for deletion SVs across all libraries. (a-c)** Distribution of sequence similarity for Hifi datasets. **(d-f)** Distribution of sequence similarity for CLR datasets. **(g-i)** Distribution of sequence similarity for ONT datasets. The evaluation was done by moderate Truvari parameters ($p$=0.5, $P$=0.5, $O$=0.01, $r$=500)

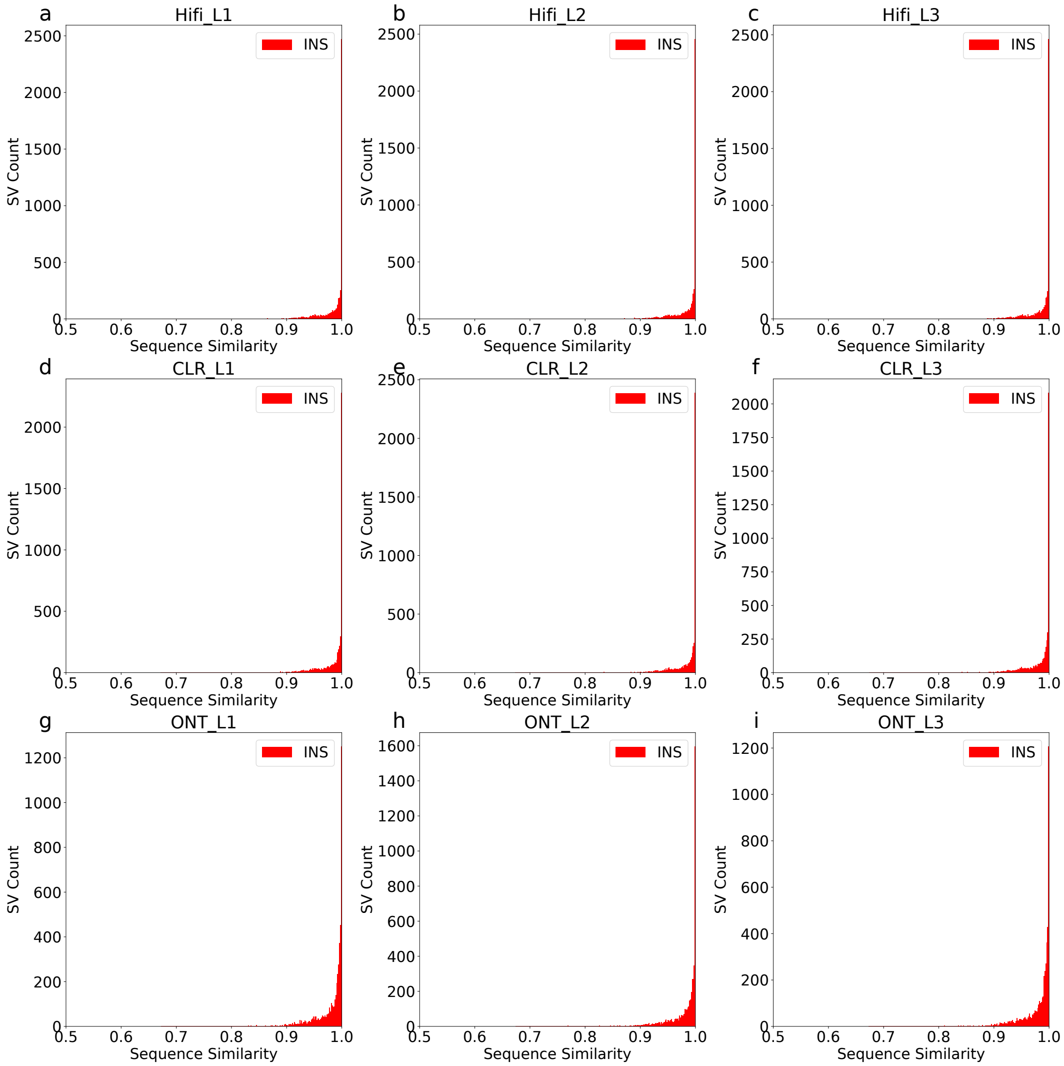

Supplementary Figure 4: **Distribution of sequence similarity for insertion SVs across all libraries. (a-c)** Distribution of sequence similarity for Hifi datasets. **(d-f)** Distribution of sequence similarity for CLR datasets. **(g-i)** Distribution of sequence similarity for ONT datasets. The evaluation was done by moderate Truvari parameters ($p$=0.5, $P$=0.5, $O$=0.01, $r$=500)

#### Supplementary References
